# Dynactin p150 promotes processive motility of DDB complexes by minimizing diffusional behavior of dynein

**DOI:** 10.1101/764761

**Authors:** Qingzhou Feng, Allison M. Gicking, William O. Hancock

## Abstract

Cytoplasmic dynein is activated by forming a complex with dynactin and the adaptor protein BicD2. We used Interferometric Scattering (iSCAT) microscopy to track dynein-dynactin-BicD2 (DDB) complexes *in vitro* and developed a regression-based algorithm to classify switching between processive, diffusive and stuck motility states. We find that DDB spends 65% of its time undergoing processive stepping, 4% undergoing 1D diffusion, and the remaining time transiently stuck to the microtubule. Although the p150 subunit was previously shown to enable dynactin diffusion along microtubules, blocking p150 enhanced the proportion of time DDB diffused and reduced the time DDB processively walked. Thus, DDB diffusive behavior most likely results from dynein switching into an inactive (diffusive) state, rather than p150 tethering the complex to the microtubule. DDB - kinesin-1 complexes, formed using a DNA adapter, moved slowly and persistently, and blocking p150 led to a 70 nm/s plus-end shift in the average velocity, in quantitative agreement with the increase in diffusivity seen in isolated DDB. The data suggest a DDB activation model in which engagement of dynactin p150 with the microtubule promotes dynein processivity, serves as an allosteric activator of dynein, and enhances processive minus-end motility during intracellular bidirectional transport.

**TOC Highlight:** Dynein-dynactin-BicD2 (DDB) is highly processive, but also shows transient pausing and diffusion, which we analyzed using iSCAT microscopy. Blocking dynactin p150 results in more diffusion of isolated DDB and a plus-end shift of kinesin-1 – DDB complexes. Thus, we conclude that p150 is an allosteric activator of dynein in the DDB complex.

## Introduction

Intracellular transport is carried out by kinesin and cytoplasmic dynein motors that walk in opposite directions along microtubules, allowing for efficient bidirectional movement of cargo^1–3^. Most cellular cargos have both kinesin motors and dynein motors bound to them ^4,5^, suggesting that robust coordination between, and regulation of, the opposite-polarity motors is required for transport; however, the underlying mechanisms are not clear. The currently prevailing model is the tug-of-war ^5,6^, in which ensembles of oppositely-directed kinesins and dyneins compete, and the stronger motor team determines the directionality. However, a number of studies have found that inhibition of one type of motor diminishes transport in both directions^7–10^; a result that suggests codependence of kinesin and dynein, and which contradicts the tug-of-war model. The tug-of-war model also does not properly account for the growing evidence that motor activity can be regulated via binding partners, and post-translational modifications of the microtubule tracks^11,12^. A more complete picture of intracellular transport must include the mechanisms by which kinesin and dynein coordinate their antagonistic activities. However, understanding this coordination first requires a more precise characterization of the individual motors, and how their activities are regulated.

Due to its diverse cellular functions, cytoplasmic dynein is known to be regulated through binding to a wide array of cargo adapter proteins^13^, a confounding factor in the effort to understand its motility. In contrast to its counterpart in yeast, it was recently discovered that mammalian dynein requires activating adapter proteins to achieve robust motility and substantial force generation *in vitro* ^14,15^. Isolated dynein adopts an inhibited phi state in which one motor domain is rotated 180 degrees with respect to the other and the two microtubule binding domain stalks are crossed, preventing microtubule binding and motility ^16^. Structural studies show that, when bound to its cofactor dynactin and the cargo adaptor BicD2, the dynein motor domains are released from the phi state and exist in an “open” conformation where they are either in a “parallel” arrangement optimal for processive walking, or in an “inverted” arrangement that allows microtubule binding but poor motility^16^. BicD2 is a coiled-coil homodimer that strengthens the normally weak interaction between the dynein tail and the dynactin filament, constraining the orientation of the dynein heads, and most likely stabilizing the parallel conformation^17–19^. This idea is supported by single molecule assays, where DDB complexes shows robust landing activity, superprocessivity, and considerably higher stall forces than dynein-dynactin or dynein alone^12,20,21^. However, a molecular description of how BicD and related adapters such as BicDR, Hook3 and Spindly work together with dynactin to activate dynein is still being resolved^12,20,22^.

A notable characteristic of activated dynein complexes *in vitro* is the broad distribution of measured velocities ^23,24^. As less than half of DDB complexes were observed to be in the activated open parallel conformation by CryoEM^16^, one explanation for this heterogeneity is that the motors switch between active and inactive states on a timescale faster than the experimental time resolution. This switching could produce periods of processive stepping interspersed with periods of pausing or 1D diffusion with zero net speed; thus, the overall speed would reflect the fraction of time the motor spends in an activated state. But what could cause this switch? One candidate is the dynactin p150 subunit, which contains a flexible linker terminating in a positively-charged CAP-Gly domain that can interact with the microtubule and is known to affect dynein motility^25^. However, the mechanism underlying this dynein velocity heterogeneity has never been investigated due to a lack of high-resolution motility data and appropriate analysis tools to objectively separate the different motility states.

Here, we apply high-resolution particle tracking and a novel switch point detection algorithm to investigate the mechanism of dynein activation by BicD2 and dynactin. Consistent with previous observations ^12,19,20,26^, DDB transitions between processive, diffusive, and stuck states. The stuck and diffusive episodes could be entirely due to p150-microtubule interactions; alternatively, they could reflect dynein being in an inhibited state that retains microtubule binding. We explored these two possibilities using a p150 antibody, previously shown to inhibit p150 interaction with microtubules^25,27–29^. We found that blocking p150 led to longer and more frequent diffusive episodes and shorter processive episodes, suggesting that the diffusive behavior of DDB results from the dynein heads rather than from p150. When DDB was complexed with kinesin-1 using a DNA adapter, blocking p150 led to a plus-ended shift in the mean velocities, in quantitative agreement with the switching behavior of DDB alone. Thus, we conclude that dynactin subunit p150 acts as an allosteric activator of dynein that accelerates switching from, and helps prevent a return to, its inhibited state.

## Methods

### 1. Plasmid constructs and DDB purification

BicD2 (25-400 a.a.)^20^ was inserted to the pET28a plasmid with an N-term StrepII tag and a C-term eGFP and His6 tag, expressed in *E. coli*, and purified by Ni column chromatography. Bovine brains were sliced and flash-frozen on dry ice at the slaughterhouse, and stored at −80 °C. To purify DDB, brain was mixed with equal volume of 50H50P buffer (50 mM Hepes, 50 mM Pipes, 2 mM MgSO_4_, 1 mM EDTA, pH 7.0), incubated in 37 °C water bath, and then homogenized in a blender, following published protocols ^20^. The lysate was clarified by centrifugation at 30,000 x g for 30 min, and the supe was mixed with equal volume A buffer (30 mM Hepes, 1 mM EGTA, 50 mM K-acetate, 2 mM Mg-acetate, 10% glycerol, pH 7.4) supplemented with 3 mM DTT, 1 mM PMSF and 0.1% NP-40 alternative^20^. The mixture was further centrifuged at 100,000 x g for 20 min, and the supe mixed with 100 nM BicD2 and incubated at 4°C for 2 hr. A column containing 2 ml of StrepTactin beads (IBA, Lifesciences) was rinsed with 3 column volumes of A buffer, the sample was applied to the column, the column was washed with A buffer, and the protein was eluted with A buffer containing 3 mM DTT and 5 mM d-Desthiobiotin (Sigma-Aldrich). The elution was used directly in single molecule experiments or flash frozen on liquid N_2_ and stored at −80 °C.

### 2. Nanoparticle functionalization of DDB

DDB containing a C-terminal GFP was attached to streptavidin-functionalized nanoparticles through a biotinylated GFP binding protein nanobody (GBP) ^30,31^. Following a previous approach^32^, a coexpression plasmid containing the BirA enzyme was constructed by inserting the GBP ^31^ sequence followed by a C-terminal Avi-tag (GLNDIFEAQKIEWH)^32^ and His_6_ tag. Biotinylated GBP was bacterially expressed and purified by Ni column chromatography. In all experiments, cover slips were washed thrice each with 70% ethanol and ddH_2_O. Microtubules were bound to the coverglass of flow cells using full-length rigor kinesin-1^32^. For landing experiments, DDB complexes were first mixed 1:1 with GBP and incubate for 5 min and then diluted to 10 nM with motility buffer, mixed with 10 nM quantum dots (incubate for 5 min), and added to the flow cell in the presence of 1 mM ATP. In Apo-lock experiments, 10 nM DDB complexes (based on GFP fluorescence) were first added to the flow cell in the absence of ATP and incubate for 5 mins to allow binding to the microtubules. After a wash to remove unbound complexes, a 10 nM solution of GBP was injected and incubated for 5 min to allow binding to BicD-GFP. Next, 10 nM streptavidin-coated quantum dots (655 nm emission; Life Technologies) were injected into the flow cell and allowed 5 min to bind to the biotinylated GBP. Finally, motility buffer containing 1 mM ATP was injected to initiate motility, and the flow cell transferred to the microscope.

### 3. Fluorescence microscopy and particle tracking

Single molecule quantum dot experiments were carried out by TIRF microscopy, as previously described^31^. 500-frame movies were taken at 20 frames/s, starting 5 mins after injecting the final motility solution, and at least 5 independent flow cells were studied for each measurement. For each field, an image was taken of the Cy5-labeled microtubules. For Ab_p150_ experiments, DDB was mixed with 25 ug/ml Ab_p150_ (BD, Biosciences, No. 610474), incubated for 30 min on ice^33^, and all subsequent solutions introduced into the flow cell also contained 25 ug/ml Ab_p150_. Image processing and kymograph analysis were performed in Image J (National Institutes of Health, Bethesda, MD). Landing rates were calculated by counting all events on a given microtubule for 10 seconds video length, and normalizing to counts to per min per microtubule length. Minimum event duration was 200 ms.

### 4. ISCAT microscopy and image processing

Flow cells for iSCAT microscopy were prepared similarly to TIRFM, with minor modifications. After Apo-lock of DDB to microtubules, 1 nM GBP was introduced and incubated for 5 min, followed by introduction of 1 nM of 30 nm gold nanoparticles (BBI Solutions) and a 5 min incubation to allow binding. Finally, ATP motility buffer was introduced and incubated for 5 min to initiate movement, the flow cell was then transferred to the microscope. The iSCAT microscope used in the work was described previously ^34^. Images were taken using custom written LabVIEW software. The videos were taken at 100 fps for 1000 frames with an effective pixel size of 32 nm. Even illumination was achieved through flat fielding before image acquisition^32^. A background image of stationary microtubules before or after particle binding was subtracted from the stack of iSCAT images, and the resulting movies were then inverted to obtain a bright gold signal on a dark background. Particle positions over time were tracked by FIESTA^35^; if no particle position was determined for 10 consecutive frames due to low signal/noise, the trace was terminated. Details for the switch detection algorithm are provided in Supplementary Information.

### 5. Kinesin-1/DDB origami experiments

DDB and *Drosophila* kinesin-1 motors (truncated at residue 560 and C-terminal GFP tagged^36^)were linked to a dsDNA scaffold following a previously published protocol employing GBP functionalized with specific DNA ^31^. To generate motors functionalized with different oligonucleotides, DDB was incubated for 15 min on ice with GBP1 in excess, and kinesin incubated with GBP2 in excess. Next, DDB-GBP1 was incubated for 15 min on ice with an excess concentration of DNA scaffold containing single-stranded overhangs on both ends and biotin on one end (**Fig. 8A**; scaffold described previously^31^). The mixture was then introduced into a flow cell containing surface-immobilized microtubules, and incubated for 5 min in the absence of ATP to allow binding of the DDB-GBP1-DNA complexes to the microtubules. The flow cell was then washed twice with A buffer containing 0.2 mg/ml casein and 10 μM Taxol to remove any unbound motors, BicD2, and GBP1, leaving only DDB with attached DNA scaffolds bound to the microtubules. An excess of kinesin-1 - GBP2 was then introduced into the flow cell and incubated for 5 min to populate the second end of the DNA scaffolds with kinesin motors. 1 nM quantum dots (633 nm emission) were then introduced into to the flow cell in the presence of ATP to label the DNA scaffolds and initiate movement, and videos were taken immediately. To determine microtubule polarity, we observed the plus-end streaming of the free GFP-labeled kinesin-1 motors in the GFP channel (**Fig. 8B**, **Supplementary Video 1**).

## Results

### Purified DDB complexes display diverse motility behavior

DDB complexes were purified from bovine brain lysate by adding recombinant BicD2, binding the complexes to StrepTactin beads (IBA Lifesciences), and eluting from the beads with d-Desthiobiotin (Sigma-Aldrich) ^37^ **(Fig. 1A, B**). The purified DDB contained a C-terminal GFP on BicD2 for visualization, but for enhanced spatiotemporal resolution, we attached streptavidin-functionalized quantum dots (Qdots) through a biotinylated GFP binding protein (GBP) nanobody ^30^ (**Fig. 1C**; see Methods for details). Using total internal reflection fluorescence (TIRF) microscopy with 50 ms exposure time, we tracked the motility of single DDB complexes along surface-immobilized microtubules and compared them to kinesin-1. Whereas kinesin-1 displayed runs with uninterrupted motility, DDB displayed three different motility behaviors: processive runs, diffusional episodes and stuck segments where no movements were detected (**Fig. 1D**). These behaviors have been observed in published DDB traces, but studies to date have generally focused only on segments of processive motility ^12,20^.

**Figure 1.**
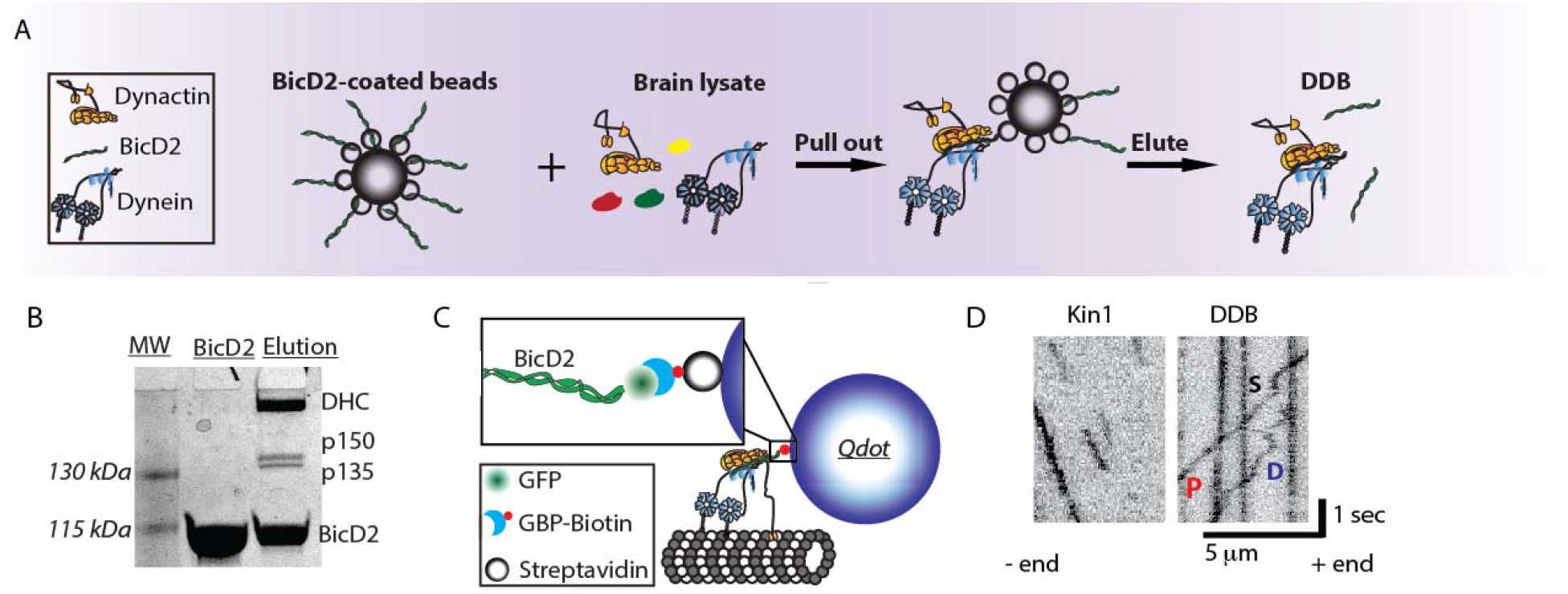
Purified DDB complex demonstrate processive, diffusive and stuck behaviors. **(A)** Schematic of DDB purification using BicD2-coated StrepTactin beads to pull dynein/dynactin from brain lysate. DDB was then eluted off of the bead. **(B)** SDS-PAGE gel of recombinant BicD2 and final purified DDB complex showing dominant bands of dynein heavy chain (DHC), dynactin components p150 and p135, and BicD2. **(C)** Tagging DDB for single molecule tracking. Biotinylated GFP binding protein (GBP) is used to link C-terminal GFP on BicD2 to streptavidin-coated quantum dots for TIRF experiments or streptavidin-coated 30-nm gold nanoparticles for iSCAT experiments. **(D)** Kymograph of kinesin-1 (left) and DDB (right) single molecule motility. DDB displays processive runs (P), diffusive episodes (D), and stuck events (S).

### Blocking dynactin p150 alters DDB landing and motility

The role of the dynactin p150 subunit in dynein activation has not been investigated, although p150 has been shown to act as both a tether and a brake in dynein-dynactin complexes^25^. To characterize how dynactin p150 alters DDB function, we utilized a p150 antibody (Ab_p150_) that has previously been shown to block the interaction of p150 with microtubules^25,27,28,33^, and compared the DDB motility in the absence and presence of Ab_p150_. We first asked what role p150 plays in the initial landing of DDB to the microtubule. Based on its tethering activity, it could enhance landing by making first contact with the microtubule and allowing the dynein heads to bind; alternatively, the runs could be all initiated by dynein heads binding (**Fig. 2A**). In the presence of antibody, the DDB landing frequency decreased by roughly three-fold compared to control (**Fig. 2B, C**). This result, consistent with previous observations ^25,38^, suggests that the initial encounter of DDB with the microtubule usually occurs through p150, although more complex mechanisms are possible. Because our DDB preparation contained a sub-fraction of p135 (**Fig 1B**), an isoform that lacks the CAP-Gly domain, it is possible that a fraction of the remaining landing events in the presence of p150 antibody represent complexes containing p135 rather than p150, meaning that our measurements provide a lower bound of the antibody effect.

**Figure 2:**
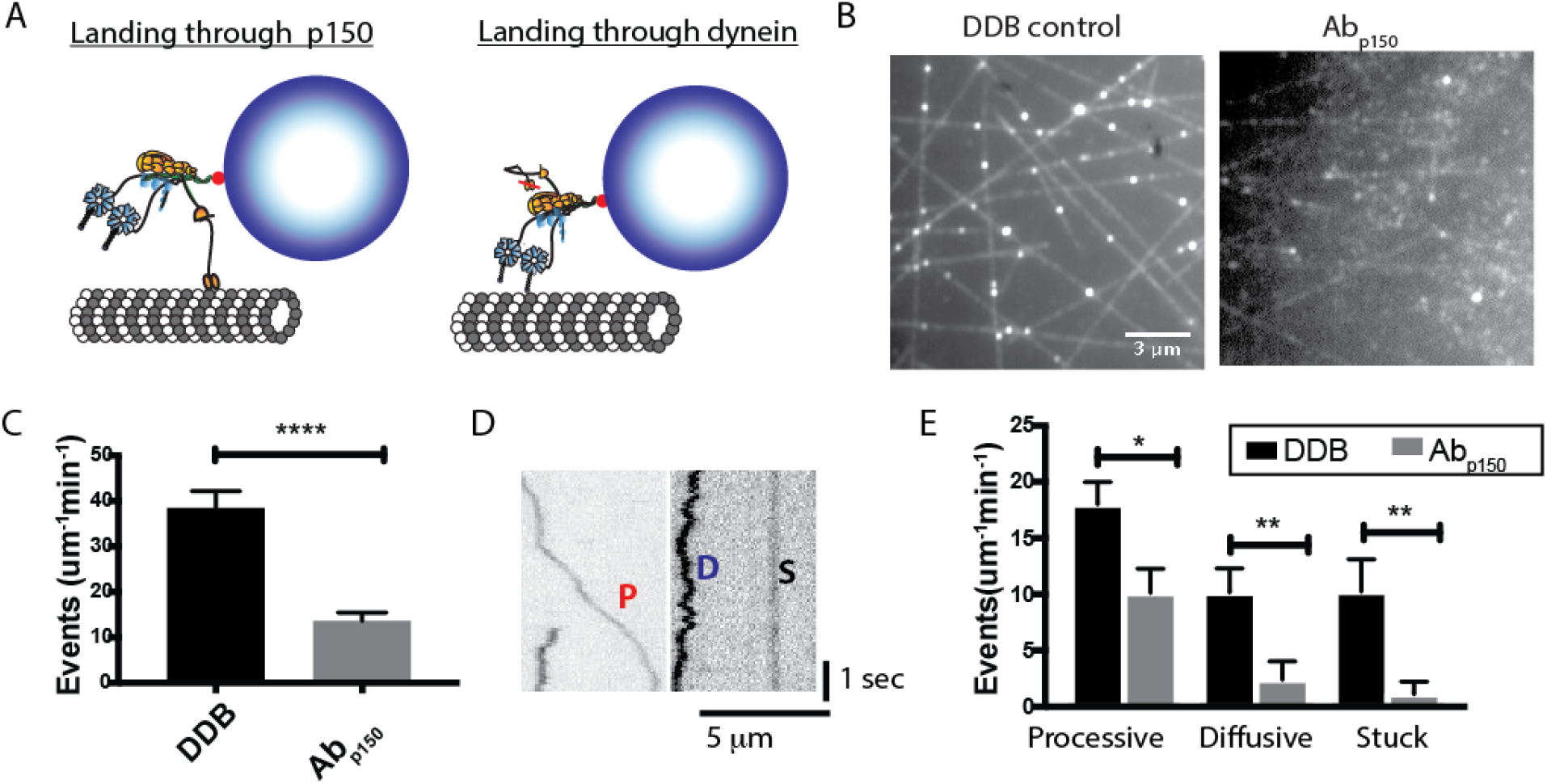
p150 of dynactin promotes landing of DDB complexes. **(A)** Diagram of landing experiment. Initial landing of Qdot-labeled DDB complexes on microtubules can occur either through dynein or through dynactin p150. **(B)** Field of microtubules and attached DDB complexes for control (left) and in the presence of Ab_p150_ (right). **(C)** Frequency of landing events in control (black, n = 10 microtubules in 50 s movie length) and Ab_p150_ (gray, n = 10 microtubules in 50 s movie length). Error bars are SEM; *** p<0.001 by t-test. **(D)** Kymographs of DDB landing events, showing processive (P), diffusive (D) and stuck (S) events. **(E)** Frequency of processive, diffusive and stuck landing events for control DDB and DDB in the presence of Ab_p150_. Error bars are SEM; *p<0.05 and **p<0.01 by t-test.

We next asked how, following initial landing of DDB on the microtubule, p150 influences dynein motility. To analyze dynein motility, the observed landing events were separated into three classes: stuck (S) complexes moved less than 100 nm overall; diffusive (D) complexes moved bidirectionally more than 100 nm for both directions with no observed unidirectional processive segments longer than 350 nm; and processive (P) complexes contained at least one segment of unidirectional movement longer than 350 nm (**Fig. 2D**). For control DDB, roughly half of the complexes that landed displayed processive motility, and the rest were split between diffusive and stuck (**Fig 2E**). Blocking dynactin p150 with the antibody reduced the frequency of processive molecules by half, and reduced the number of diffusional and stuck complexes to near zero (**Fig. 2E**). A simple interpretation of the drop in processive events is that half of these events occur when dynein initially contacts the microtubule and the other half when dynactin p150 initially contacts the microtubule. It follows that molecules that solely diffuse along or stick to the microtubule without any processive behavior initially bind to the microtubule through their dynactin p150 subunit, and their dynein is either in an inhibited state or possibly damaged.

### Dynactin p150 enhances processive and diminishes diffusive behavior of DDB

To select for active DDB complexes, we introduced DDB into the chamber in the absence of ATP, such that active dynein bound to the immobilized microtubules in the apo (no nucleotide) state. Following this “Apo-lock”, any unbound complexes were washed out with nucleotide-free buffer, and movement was initiated by flowing ATP containing buffer into the chamber (**Fig. 3A**). Here “active DDB complexes” are defined as those that bind microtubules statically in the apo state and release in the ATP state. As with the landing experiments, processive, diffusive, and stuck behaviors were all observed (**Fig. 3B**). In the absence of dynactin p150 antibody, roughly half of the complexes moved processively upon ATP addition, whereas the other half either remained stuck in ATP (~40%) or displayed only diffusive behavior (~10%) (**Fig 3C**; **DDB**). In the presence of dynactin p150 antibody, the fraction of processive complexes fell, while the fraction of diffusive complexes increased (**Fig 3C**; **p150**). This is the opposite of what would be predicted if p150 were simply acting as a diffusional tether; if that were the case, there should be fewer diffusive complexes when p150 is blocked. Although informative, this analysis categorized every particle as processive, diffusive, or stuck, which is relatively coarse. Deeper understanding of how dynein is activated in the DDB complex and how dynactin p150 contributes to this activation requires a more detailed analysis of the processive complexes, where DDB switches between processive, diffusive and stuck states within a single run.

**Figure 3.**
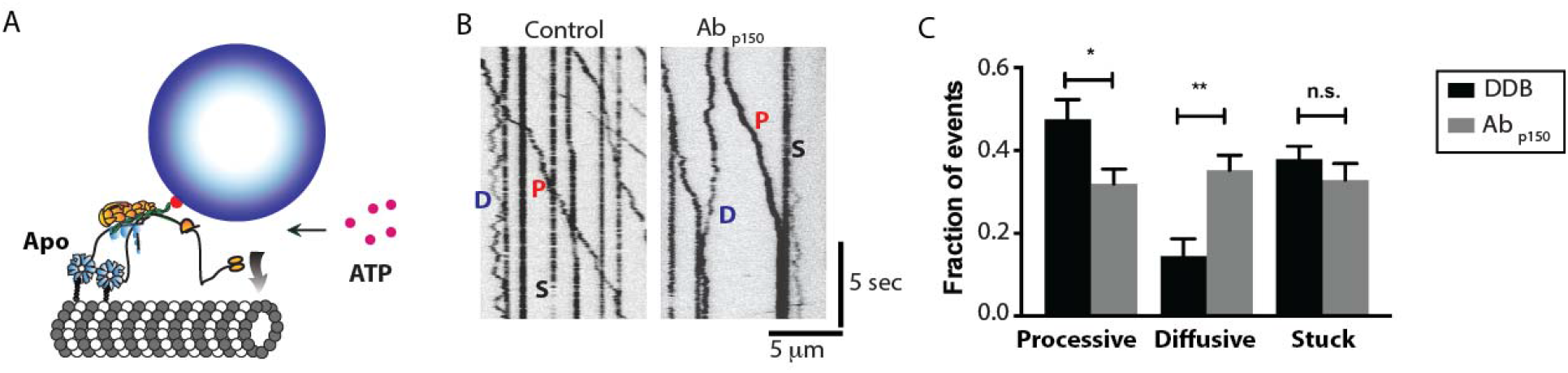
Blocking p150 dynactin leads to fewer processive events and more diffusive events. **(A)** Diagram of the Apo-lock experiment. DDB complexes bind to immobilized microtubules in absence of ATP, and ATP buffer is flushed into the system to initiate motility. **(B)** Kymograph of DDB motility 5 min after flowing in ATP buffer for control (left) and in presence of Ab_p150_ (right). Processive (P), diffusive (D) and stuck (S) events are noted. **(C)** Average fraction of processive, diffusive and stuck traces across 10 kymographs for control (black) and Ab_p150_ group (gray). Error bars are SEM; * denotes p<0.05 (t-test); n.s., not significantly different.

### p150 promotes switching into and prevents switching out of the processive state

To investigate how p150 affects the kinetics of DDB switching between different motility states, we enhanced our temporal resolution by attaching 30 nm gold nanoparticles to BicD2 in our DDB complex and tracking them with Interferometric Scattering (iSCAT) microscopy. An iSCAT image is formed by interference between light scattered by the gold particle and light reflected at the glass-water interface of the sample (**Fig. 4A**)^39^. With this approach, unlabeled microtubules and gold particles can be visualized simultaneously, with particles appearing as dark spots on a bright background (**Fig. 4B**). After subtracting an image of the stationary microtubule and inverting the image to produce a bright particle on a dark background, the point-spread function (PSF) of the gold particle can be fit by a 2-D Gaussian distribution (**Fig. 4D**) to achieve nm-scale spatial precision. By analyzing movies with FIESTA software ^35^, x-y position over time data was collected at 100 frames/s, which we found to be the optimal temporal resolution for this work.

**Figure 4.**
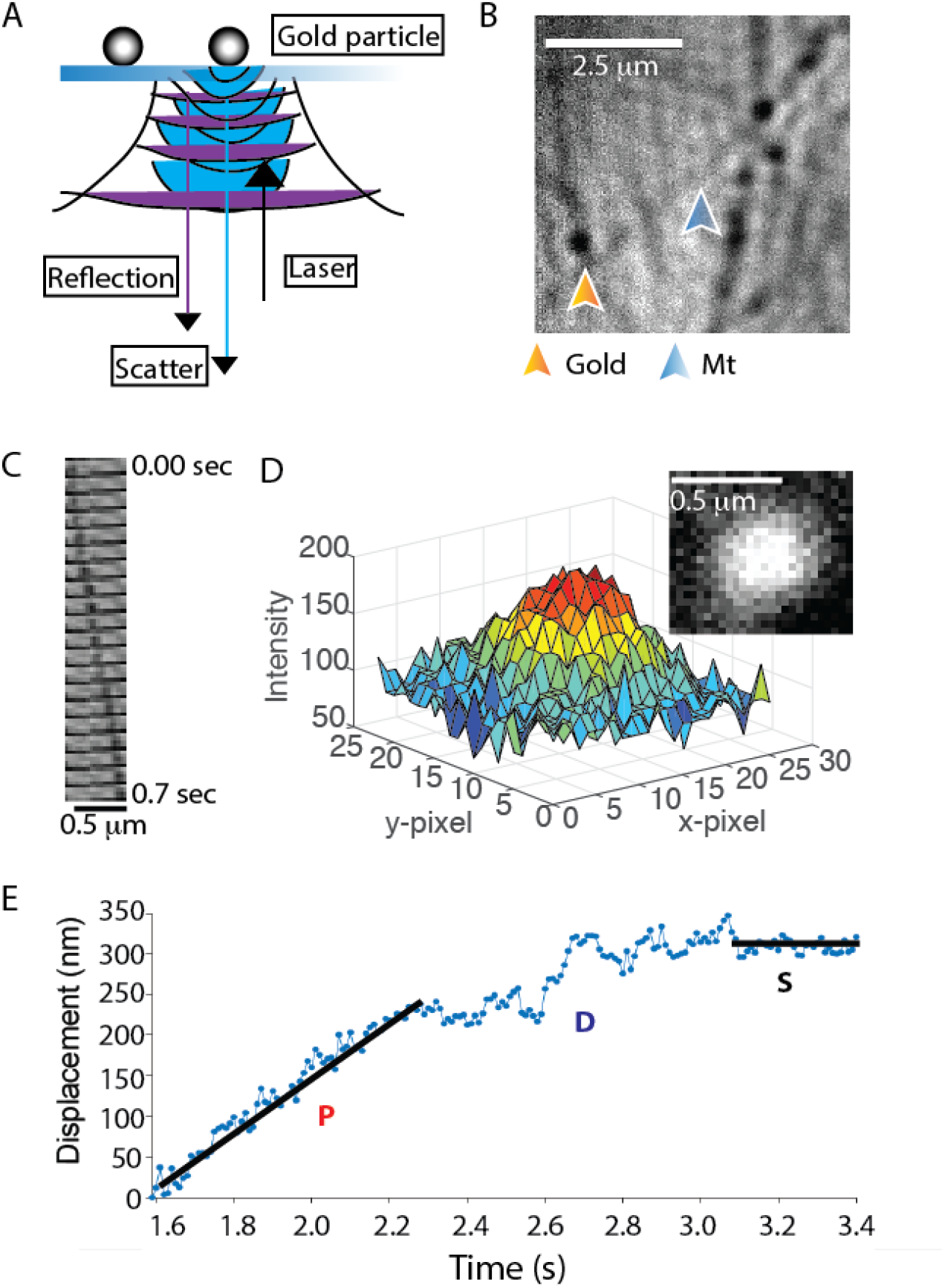
Single molecule DDB tracking by iSCAT microscopy. **(A)** Diagram of iSCAT microscopy. Image is formed by scattered light from the gold nanoparticle (blue) interfering with reflected light from the glass-water interface (purple). **(B)** iSCAT image of a field of gold nanoparticle-labeled DDB bound to surface-immobilized microtubules. Image shown is generated from a raw image by flat fielding, which corrects inhomogeneous illumination across the field. **(C)** Montage of a gold particle-labeled DDB moving along an immobilized microtubule; each image is 35 msec apart. **(D)** Plot of pixel intensity of a gold nanoparticle (image in inset), which is fit by a 2-D Gaussian for sub-pixel localization. Image is generated by subtracting image of the stationary microtubule (taken later in the movie when no gold-labeled motor is present) and inverting image to obtain bright particle on dark background. See also **Supplementary Movie S1**. **(E)** Distance vs time trace of a single DDB, demonstrating processive (P), diffusive (D), and stuck (S) episodes in the same trace. Lines represent linear regressions to hand-selected segments.

By processing the traces to obtain linear distance along the microtubule over time, DDB complexes clearly switch between processive, diffusive and stuck states during a given trace (**Fig. 4E**). Although some phases such as long processive or stuck phases are readily identifiable, diffusive phases are particularly difficult to define, despite the high spatiotemporal resolution. Thus, we developed an objective algorithm for classifying processive, diffusive and stuck durations within a single trace. The algorithm, described fully in Supplementary Methods, uses a 10-frame running window and calculates the positional standard deviation, the slope, and the residual around the slope for each point in the trace. Based on defined cutoff values that are optimized with simulations, each point is classified and the traces are then broken into continuous segments of at least 100 msec (10 frame) duration each. A gallery of processed traces is shown in **Fig. 5**, with colors indicating processive (red), diffusive (blue), and stuck (black) states.

**Figure 5.**
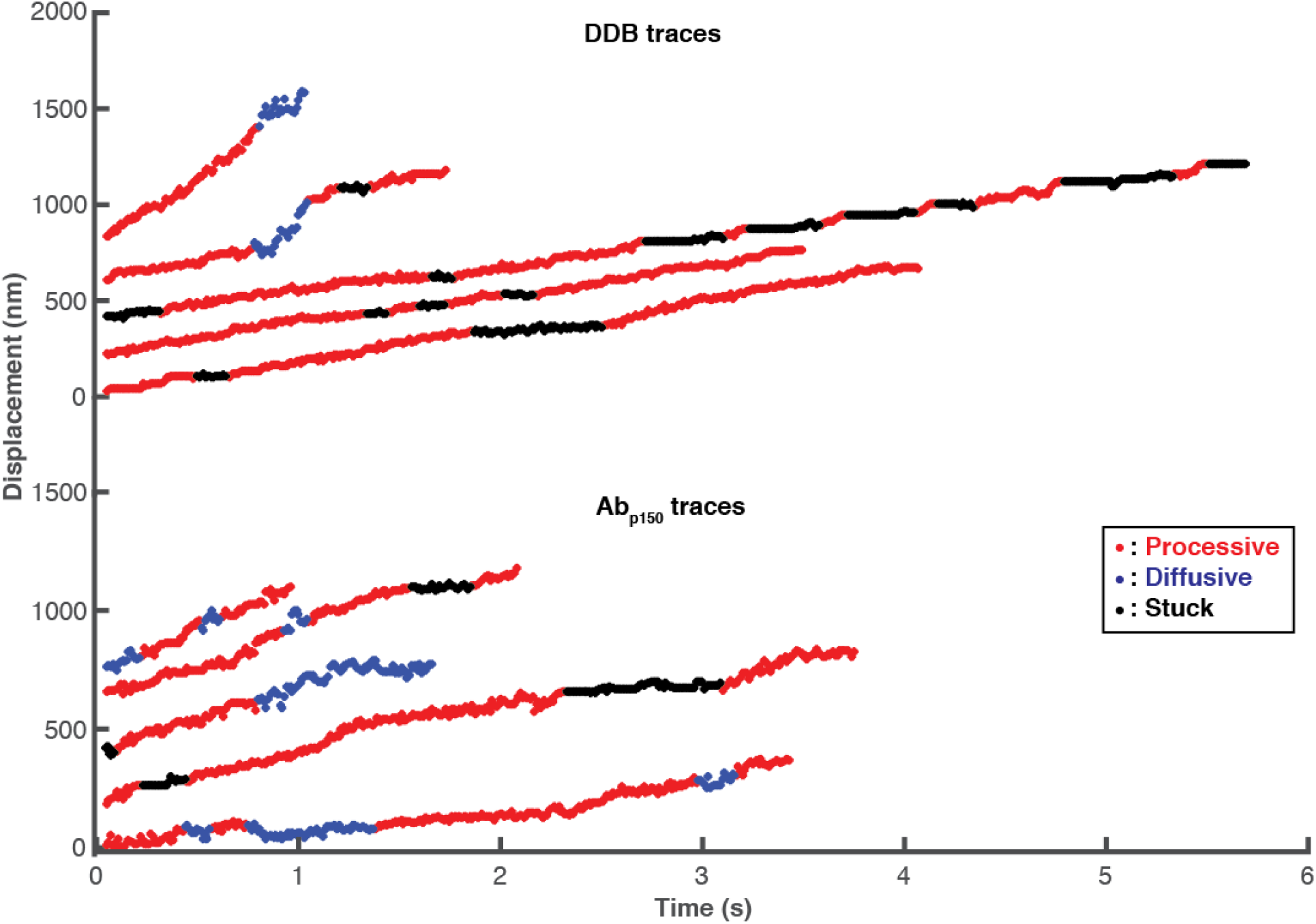
High-resolution DDB tracking and motility state identification. Sample traces of control DDB (top) and DDB in presence of Ab_p150_ (bottom) taken at 100 frames/s by iSCAT microscopy and processed with the state switching algorithm. Processive segments are labeled in red, diffusive episodes in blue, and stuck durations in black.

Dividing each single molecule trajectory into different phases, or motility states, provides distributions of time the motor spends in each state, as well as the switching rates between the three states. For DDB under control conditions, processive segments had the longest duration at 0.81 s, followed by stuck (0.53 s) and diffusive (0.23 s) phases (**Fig. 6 A**). The most frequent switching was between stuck and processive states (**Fig. 6 A inset**), meaning that there were relatively frequent short pauses during processive stepping. The second most common switching was between processive and diffusive states. These two behaviors can be seen qualitatively in **Fig. 5** as short black and blue phases interspersed in the relatively long processive runs in red.

**Figure 6.**
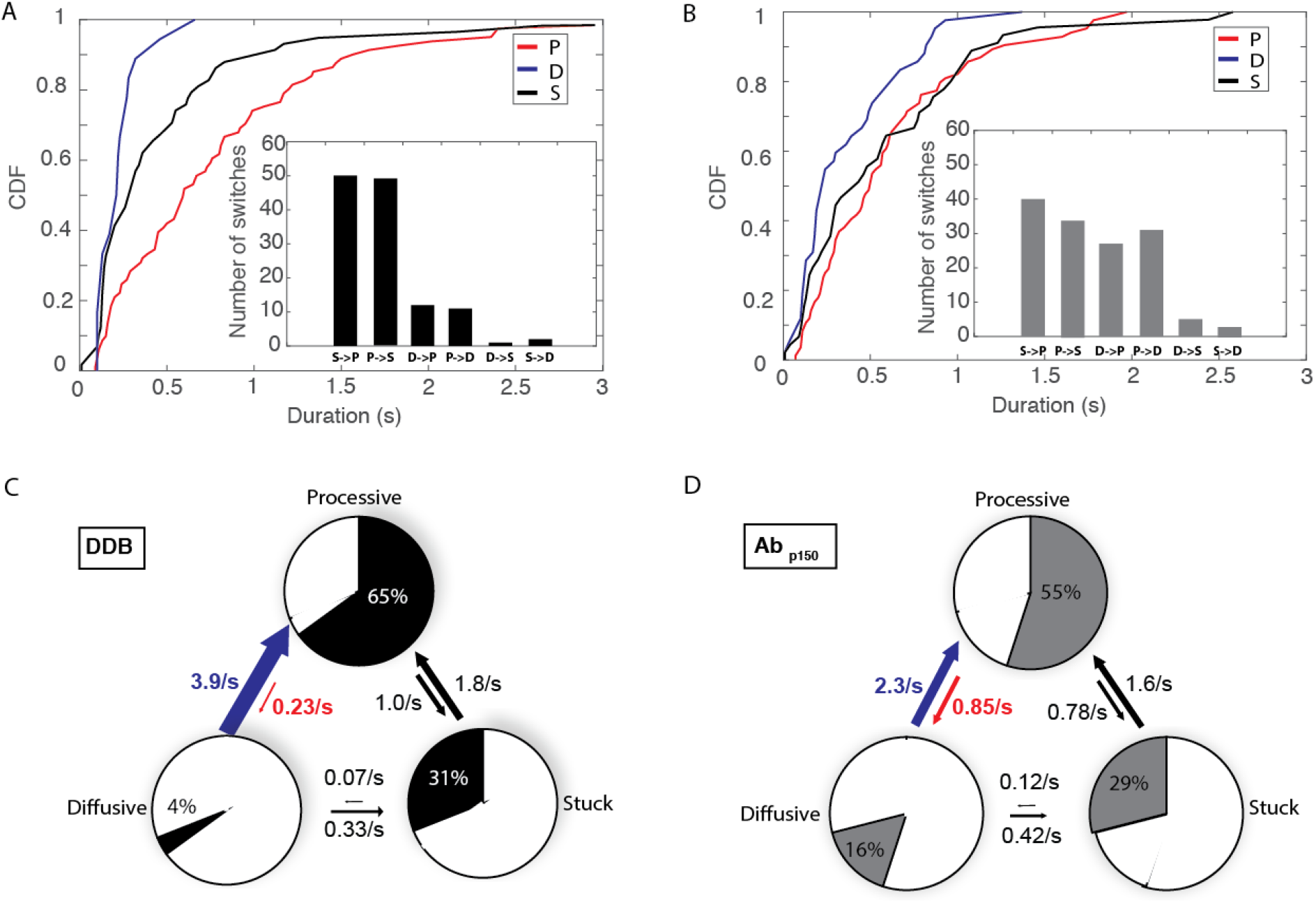
p150 shortens diffusive segments and elongates processive segments. **(A)** Cumulative distributions of processive, diffusive and stuck segment durations for control DDB. Mean durations were 0.81 s for processive, 0.23 s for diffusive, and 0.53 s for stuck states. Inset: Number of detected state switches over 93 s total analyzed time from 31 molecules. **(B)** Cumulative distributions of processive, diffusive and stuck segment durations for DDB in the presence of Ab_p150_. Mean durations were 0.61 s for processive, 0.37 s for diffusive, and 0.60 s for stuck states. Inset: Number of detected state switches for Ab_p150_ group over 100 s total analyzed time from 32 molecules. **(C)** State switching diagram showing first-order switching rates between states and fraction of time spent in each state for control DDB. Blue-colored arrow denotes the most significant decrease in switching rate with Ab_p150_, while red arrow denotes the most significant increase in switching rate. **(D)** State switching diagram showing first-order switching rates between states and fraction of time spent in each state DDB in the presence of Ab_p150_.

From the state durations and switching frequencies, we created a kinetic model for how DDB switches between processive, diffusive and stuck states and what fraction of the time the motors spend in each state. Each state (P, D and S) has two transitions in and two transitions out, and all transitions were assumed to be first order based on the roughly exponential distribution profiles in **Fig. 6A**. The transition rate out of any given state equals the sum of the two rate constants exiting that state, and the relative rates between the two exit paths are taken from the measured switch rates in **Fig. 6A inset**. The switching model (**Fig. 6C**) provides a wealth of information. First, the motors spend 65% of the time in the processive state and most of the remaining time (31%) in the stuck (paused) state. Second, if the motors ever enter the diffusive state or the stuck state, they rapidly transition back to the processive state (at 3.9 s^−1^ and 1.8 s^−1^, respectively). Finally, transient events that break up the processive runs are more often short pauses (occurring at a frequency of 1 s^−1^), rather than diffusive episodes (at a frequency of 0.23 s^−1^).

To understand the role of dynactin p150 in dynein activation and diffusional tethering, we repeated the analysis for DDB in the presence of the p150 antibody. When dynactin p150 was blocked, the duration of the processive segments decreased to 0.61 s, while the duration of diffusional segments increased to 0.37 s (**Fig. 6B**). Compared to control, switching occurred less frequently between processive and stuck states, and more frequently between processive and diffusive states (**Fig. 6B inset**). As clearly shown in the kinetic model (**Fig. 6D**), blocking p150 caused the motor to spend less time in the processive state (55%) and more time in the diffusive state (16 %). The kinetic explanation for this (highlighted by red and blue arrows in **Fig. 6 C and D**) is that the presence of p150 causes DDB to switch 69% *more* frequently from the diffusive state into the processive state and to switch 73% *less* frequently out of the processive state back to the diffusive state. A structural interpretation of these results is shown in **Fig. S7** and discussed more fully below. To conclude, allowing p150 to interact with the microtubule both promotes and stabilizes the processive state of dynein in the DDB complex.

### p150 enhances minus-end directionality of kinesin-DDB complexes

Based on the finding that p150 enhances the time DDB spends in the processive state, it follows that p150 should enhance dynein’s ability to compete against kinesin-1 in a tug-of-war such as occurs during intracellular bidirectional transport. To investigate this possibility, we reconstituted the kinesin-dynein bidirectional transport system *in vitro* using a DNA origami scaffold. One kinesin-1 motor and one DDB were connected through a DNA scaffold functionalized with a quantum dot (**Fig. 8A**), and the complexes tracked by TIRF microscopy. Consistent with previous *in vitro* tug-of-war experiments^12^, long duration events were observed with mean velocities much slower than either individual unloaded motor speed, indicating that both motors engaged with the microtubule (**Fig. 8B**). To investigate the role of p150 in bidirectional transport, we compared the mean velocities of traces in the absence and presence of Ab_p150_. The simple prediction is that, if blocking p150 increases the fraction of time the motor is in the diffusive state (from 4% to 16%; **Fig 6C, D**) then the mean velocity should shift toward the plus-end in the presence of the antibody. For the control case, we measured a mean velocity of −9.1 ± 9.2 nm/s (mean ± SEM, N = 33) toward the minus-end (**Fig. 8D**). In the presence of Ab_p150,_ the mean velocity shifted to 62 ± 17 nm/s (mean ± SEM, N = 32; **Fig. 8D**), a statistically significant change (p = 0.0004 by two-tailed t=test). In addition, the proportion of complexes with a net plus-end directionality increased from 42% in the control case to 75% when p150 was blocked (**Fig. 8E**).

The +71 ± 19 nm/s shift in the mean velocity when p150 was blocked is in good quantitative agreement with our switching model, as follows. The diffusive episodes analyzed to develop the detection algorithm had a 1D diffusion constant of D = 20,000 nm^2^/s by mean-squared displacement analysis (Fig S1D). This can be converted to a drag coefficient, γ, using D = k_B_T/γ, where Boltzman’s constant times absolute temperature, k_B_T = 4.1 pN-nm^40^. The resulting drag coefficient of γ = 0.0002 pN-s/nm means that a DDB in the diffusive state that is being pulled by a kinesin moving at v = 500 nm/s should produce a drag force (F = γ*v) of only 0.1 pN, which should not slow the kinesin^41^. From the switching model in **Fig. 6C, D**, blocking p150 increased the fraction of time in the diffusive state by 12%, from 4% to 16%. If the complexes move at 500 nm/s for 12% of the time, this would contribute 0.12 * 500 nm/s = 60 nm/s of mean plus-end velocity, which closely matches the observed +71 +19 nm/s increase. Thus, we interpret the slow kinesin-DDB transport velocities to reflect the antagonistic motors pulling against one another with DDB stochastically switching between motile states. Blocking p150 shifts DDB toward more time in the diffusive state that kinesin readily pulls against, resulting in a plus-end shift in the net transport velocity.

## Discussion

Understanding how specific intracellular cargo are targeted to their proper cellular locations requires understanding how bidirectional transport is regulated, which in turn requires understanding the regulation of dynein activation. By tracking DDB complexes at high temporal resolution and applying our change-point detection algorithm, we found that in the DDB complex, dynein switches between active and inactive states at rates exceeding 1 s^−1^ (**Fig. 6C**). This analysis leads to two questions. First, to what degree is dynactin p150 tethering the complex during processive motility? Second, do the diffusive and stuck periods reflect only p150 interacting with the microtubule, only inhibited dynein interacting with the microtubule, or some combination of the two? Blocking p150 provides the following insights. First, the observation that blocking p150 results in more, rather than fewer diffusive complexes (**Fig. 3C**) suggests that diffusive DDB behavior, also observed by others^42^, reflects complexes where dynein is in an inhibited state that binds to microtubules, rather than complexes that are tethered solely through p150. Second, the longer durations of diffusive segments following p150 block (**Fig. 6B**) suggests that switching into this state during processive runs reflects dynein switching into an inhibited state, rather than dynein detaching from the microtubule while p150 maintains overall microtubule association of the complex. Third, the finding that the switching rate into and out of the stuck state during processive runs was unaffected by Ab_p150_ (**Fig. 6 C, D)** suggests that this paused state is inherent to the stepping mechanism of dynein or at least that p150 alone is not sufficient to prevent the formation of this inhibited state. And last, there was no significant difference between mean velocities of processive segments in control versus p150 block (**Fig. S6**), arguing that p150 does not act as a brake slowing dynein in the DDB complex, contrary to previous observations on dynein-dynactin complexes lacking BicD2^25^.

Based on recent structural studies, we can make tentative structural assignments to our identified functional states of dynein. Because the dynein-dynactin-DDB structure is incompatible with dynein being in the inhibited “phi” state^16^, we interpret our DDB complexes to reflect dynein in the “open” conformation, with the heads either in an “open-parallel” configuration optimal for stepping, an “open-inverted” conformation that can bind to microtubules but not processively step (**Fig. 7A**). Similarly, we hypothesize that in the DDB structure, p150 is sterically free and able to reversibly interact with microtubules ^43^. This leads to four possible states (**Fig 7A)**, with dynein being in either an open-parallel or open-inverted conformation and p150 either interacting with the microtubule and constraining the dynactin orientation, or p150 being free and dynactin being less conformationally constrained. In this model, when p150 interacts with the microtubule, the open-parallel conformation of dynein is favored, whereas blocking p150 from binding to the microtubule biases the motor toward the open-inverted conformation (highlighted states in **Fig. 7A**).

**Figure 7.**
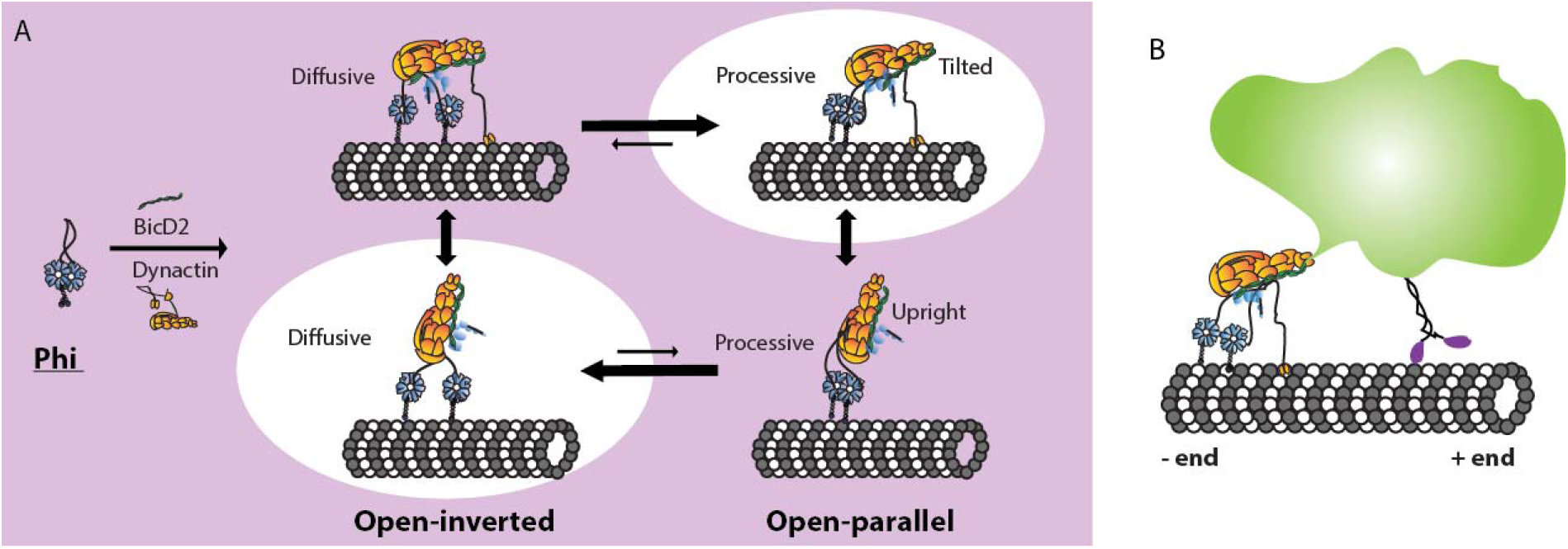
Proposed model for DDB motility enhancement by p150 dynactin. **(A)** Dynein can reside in the inactive phi conformation in solution, but forming a DDB complex results in dynein switching to an open conformation. In the open-inverted conformation, DDB is more likely to diffuse along microtubules, while in the open-parallel conformation DDB is more processive. (**Top**) p150 interaction with the microtubule promotes a tilted dynactin geometry that stabilizes the open-parallel conformation of dynein and results in enhanced processivity. (**Bottom**) Blocking p150 causes dynactin to adopt a more flexible upright geometry that promotes the open-inverted conformation of dynein and results in DDB diffusing on the microtubule. **(B)** Implications for bidirectional cargo transport in cells: enhancement of DDB processivity by p150 promotes net minus-end cargo transport.

**Figure 8:**
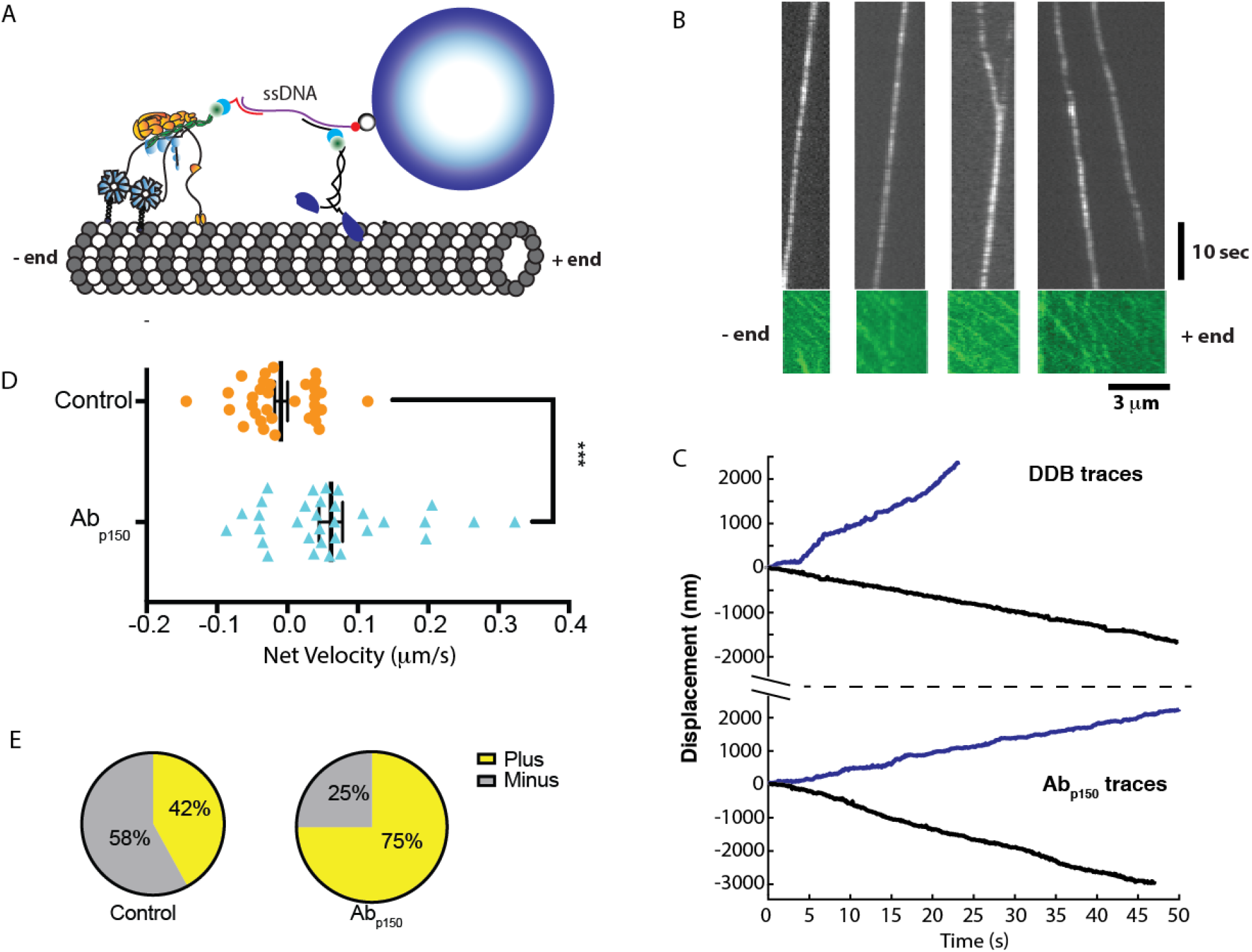
p150 activates DDB in kinesin-DDB bidirectional transport. **(A)** Diagram of the reconstituted bidirectional transport system. Single kinesin-1 and DDB are connected through ssDNA-functionalized GBP1 and GBP2 adapters to a dsDNA scaffold, linked at its biotinylated 5’ end to a streptavidin-coated Qdot. **(B)** Kymographs Qdot-labeled DDB-kinesin-1 (top) in the 647 nm channel, and the excess kinesin-1 motors streaming to the plus end in the GFP channel (bottom), used to identify the polarity of the microtubule. See also **Supplemental Movie S2**. **(C)** Sample traces of DDB-kinesin-1 for control (**top**) and Ab_p150_ group (**bottom**). **(D)** Velocities of the control group (orange; −9.1± 9.2 nm/s (mean ± SEM, n=33)) and the Ab_p150_ group (blue; 62 ±17 nm/s (mean ± SEM, n=32)) calculated by from linear regression to entire traces. The two groups were significantly different by two-tailed t-test, ***p<0.0005. **(E)** Percent of plus-end directed cargos (yellow) and minus end directed cargos (grey) for control DDB-kinesin-1 group (left) and Ab_p150_ group (right).

Instead of predominantly acting as a diffusive tether in the DDB complex, our data support a model in which p150 is an allosteric activator of dynein. The clearest evidence for this is the faster switching into the processive state and slower switching out of the processive state in the control compared to the p150 block (**Fig. 6 C, D** and **Fig. S7**). Assuming that the action of p150 is through binding to the microtubule rather than binding to the dynein heads, how could this work? Recent studies investigating the regulatory protein Lis1 and adapters like BicD2 and Hook3 that can form complexes containing two dyneins have converged on a model in which a second dynein (or even the linker and tail of a second dynein) enhances motility by stabilizing the first dynein in the open-parallel state^16,19,44,45^. Based on this, a possible explanation for p150 enhancement of motility is that when p150 is tethered to the microtubule, it orients the dynactin filament, and hence the dynein heads, in a conformation that favors the open-parallel conformation (**Fig. 7A**). Conversely, if p150 does not stabilize dynactin on the microtubule, the dynactin filament, and the two dynein heads are free to adopt multiple conformations including the non-motile open-inverted state that either diffuses along or sticks to microtubules.

In contrast to the rapid switching behavior of isolated DDB, kinesin-1 – DDB complexes displayed long duration events having slow mean velocities and both plus- and minus-end net directionalities. Work by others has also shown that adapters that more fully activate dynein generate a greater net minus-end directionality in kinesin-dynein complexes^12,44^. Because kinesin acts as an effective tether to maintain association with the microtubule in kinesin-DDB complexes, p150 is not expected to play a tethering role. However, the significant plus-end velocity shift seen upon p150 inhibition demonstrates that p150 plays an activating role even when dynein is subjected to plus-end forces from kinesin-1. Furthermore, the +71±19 nm/s shift in average speed upon p150 inhibition can be quantitatively explained by the 12% shift of DDB into the diffusive state identified by the switch point detection algorithm (**Fig. 6**). Therefore, p150 can modulate bidirectional transport in cells by enhancing dynein motility and making it a stronger opponent to kinesin-1.

Whereas kinesins achieve functional diversity through gene duplication, there is only one dynein heavy chain in the genome; thus regulation of dynein motor properties and cargo interactions must be achieved through diversity in cargo adapters and exogenous regulatory proteins^46^. Understanding dynein activation is important because during bidirectional cargo transport in cells, any regulation of dynein will alter its competition with kinesin, and hence affect cargo speed and directionality. By applying single-molecule iSCAT tracking with our novel switch-point detection algorithm, we identify switches between active and inhibited motor states in DDB and show that p150 affects the switching rates between these states. Thus, in addition to acting as a diffusional tether that can enhance dynein run lengths, p150 can enhance dynein stepping both in isolated DDB complexes and in antagonistic assemblies of DDB and kinesin-1, and as such should be added to the list of dynein activating proteins.

## Acknowledgements

We thank Richard J. McKenney for generously providing the BicD2 plasmid and advice on DDB purification, Geng-Yuan Chen for assistance with the switch-point detection algorithm, Keith J. Mickolajczyk for iSCAT microscopy mentoring, and members of the W.O.H. laboratory for helpful discussions. This work was supported by NIH R01GM121679 and R01GM122082 to W.O.H.

## Author contributions

Q.F. and W.O.H. designed research; Q.F. performed experiments and wrote the algorithm; A.M.G. and Q.F. carried out iSCAT experiments and image processing; Q.F. wrote the paper and Q.F., W.O.H. and A.M.G. edited the paper.

**The authors declare no conflict of interest**

## References

1. Hirokawa, N., Niwa, S. & Tanaka, Y. Molecular motors in neurons: Transport mechanisms and roles in brain function, development, and disease. Neuron 68, 610–638 (2010).

2. Gross, S. P. et al. Interactions and regulation of molecular motors in Xenopus melanophores. J. Cell Biol. 156, 855–865 (2002).

3. Hancock, W. O. Bidirectional cargo transport: Moving beyond tug of war. Nat. Rev. Mol. Cell Biol. 15, 615–628 (2014).

4. Ligon, L. A., Tokito, M., Finklestein, J. M., Grossman, F. E. & Holzbaur, E. L. F. A Direct Interaction between Cytoplasmic Dynein and Kinesin I May Coordinate Motor Activity. J. Biol. Chem. 279, 19201–19208 (2004).

5. Hendricks, A. G. et al. Motor Coordination via a Tug-of-War Mechanism Drives Bidirectional Vesicle Transport. Curr. Biol. 20, 697–702 (2010).

6. Müller, M. J. I., Klumpp, S. & Lipowsky, R. Tug-of-war as a cooperative mechanism for bidirectional cargo transport by molecular motors. (2008).

7. Welte, M. A., Gross, S. P., Postner, M., Block, S. M. & Wieschaus, E. F. Developmental regulation of vesicle transport in Drosophila embryos: Forces and kinetics. Cell 92, 547–557 (1998).

8. Goldberg, D. J. Microinjection into an identified axon to study the mechanism of fast axonal transport. Proc. Nati Acad. Sci. USA 79, 4818–4822 (1982).

9. Martin, M. et al. Cytoplasmic Dynein, the Dynactin Complex, and Kinesin Are Interdependent and Essential for Fast Axonal Transport. Mol. Biol. Cell 10, 3717–3728 (1999).

10. Waterman-Storer, C. M. et al. The interaction between cytoplasmic dynein and dynactin is required for fast axonal transport. Proc. Natl. Acad. Sci. 94, 12180–12185 (2002).

11. Monroy, B. Y. et al. Competition between microtubule-associated proteins directs motor transport. Nat. Commun. 9, 1–12 (2018).

12. Belyy, V. et al. The mammalian dynein/dynactin complex is a strong opponent to kinesin in a tug-of-war competition. 18, 1018–1024 (2017).

13. Olenick, M. A. & Holzbaur, E. L. F. Cell science at a glance dynein activators and adaptors at a glance. J. Cell Sci. 132, 1–7 (2019).

14. Trokter, M., Mucke, N. & Surrey, T. Reconstitution of the human cytoplasmic dynein complex. Proc. Natl. Acad. Sci. 109, 20895–20900 (2012).

15. King, S. J. & Schroer, T. A. Dynactin increases the processivity of the cytoplasmic dynein motor. Nat. Cell Biol. 2, 20–24 (2000).

16. Zhang, K. et al. Cryo-EM Reveals How Human Cytoplasmic Dynein Is Article Cryo-EM Reveals How Human Cytoplasmic Dynein Is Auto-inhibited and Activated. Cell 169, 1303–1314.e18 (2017).

17. Splinter, D. et al. BICD2, dynactin, and LIS1 cooperate in regulating dynein recruitment to cellular structures. Mol. Biol. Cell 23, 4226–4241 (2012).

18. Sladewski, T. E. et al. Recruitment of Two Dyneins to an mRNA-Dependent Bicaudal D Transport Complex. 2, (2018).

19. Urnavicius, L. et al. Cryo-EM shows how dynactin recruits two dyneins for faster movement. Nature 554, 202–206 (2018).

20. McKenney, R. J., Huynh, W., Tanenbaum, M. E., Bhabha, G. & Vale, R. D. Activation of cytoplasmic dynein motility by dynactin-cargo adapter complexes. Science 345, 337–341 (2014).

21. Schlager, M. A., Hoang, H. T., Urnavicius, L., Bullock, S. L. & Carter, A. P. In vitro reconstitution of a highly processive recombinant human dynein complex. EMBO J. 33, 1855–1868 (2014).

22. Schroeder, C. M. & Vale, R. D. Assembly and activation of dynein-dynactin by the cargo adaptor protein Hook3. J. Cell Biol. 214, 309–318 (2016).

23. Olenick, M. A., Tokito, M., Boczkowska, M., Dominguez, R. & Holzbaur, E. L. F. Hook adaptors induce unidirectional processive motility by enhancing the Dynein-Dynactin interaction. J. Biol. Chem. 291, 18239–18251 (2016).

24. Gutierrez, P. A., Ackermann, B. E., Vershinin, M. & Mckenney, R. J. Differential effects of the dynein-regulatory factor Lissencephaly-1 on processive dynein-dynactin motility. J. Biol. Chem. 292, 12245–12255 (2017).

25. Ayloo, S. et al. Dynactin functions as both a dynamic tether and brake during dynein-driven motility. Nat. Commun. 5, 1–11 (2014).

26. Grotjahn, D. A. et al. Cryo-electron tomography reveals that dynactin recruits a team of dyneins for processive motility. Nat. Struct. Mol. Biol. 25, 203–207 (2018).

27. Holzbaur, E. L. F. et al. Homology of a 150K cytoplasmic dynein-associated polypeptide with the Drosophila gene Glued. Nature 351, 579–583 (1991).

28. Waterman-Storer, C. M., Karki, S. & Holzbaur, E. L. F. The p150(Glued) component of the dynactin complex binds to both microtubules and the actin-related protein centractin (Arp-1). Proc. Natl. Acad. Sci. U. S. A. 92, 1634–1638 (1995).

29. Ross, J. L., Wallace, K., Shuman, H., Goldman, Y. E. & Holzbaur, E. L. F. Processive bidirectional motion of dynein-dynactin complexes in vitro. Nat. Cell Biol. 8, 562–570 (2006).

30. Kubala, M. H., Kovtun, O., Alexandrov, K. & Collins, B. M. Structural and thermodynamic analysis of the GFP:GFP-nanobody complex. Protein Sci. 19, 2389–2401 (2010).

31. Feng, Q., Mickolajczyk, K. J., Chen, G. Y. & Hancock, W. O. Motor Reattachment Kinetics Play a Dominant Role in Multimotor-Driven Cargo Transport. Biophys. J 114, 400–409 (2018).

32. Mickolajczyk, K. J. et al. Kinetics of nucleotide-dependent structural transitions in the kinesin-1 hydrolysis cycle. Proc. Natl. Acad. Sci. U. S. A. 112, E7186–E7193 (2015).

33. Ross, J. L., Wallace, K., Shuman, H., Goldman, Y. E. & Holzbaur, E. L. F. Processive bidirectional motion of dynein – dynactin complexes in vitro. 8, (2006).

34. Mickolajczyk, K. J., Geyer, E. A., Kim, T., Rice, L. M. & Hancock, W. O. Direct observation of individual tubulin dimers binding to growing microtubules. Proc. Natl. Acad. Sci. U. S. A. (2019). doi:10.1073/pnas.1815823116

35. Ruhnow, F., Zwicker, D. & Diez, S. Tracking single particles and elongated filaments with nanometer precision. Biophys. J 100, 2820–2828 (2011).

36. Shastry, S. & Hancock, W. O. Interhead tension determines processivity across diverse N-terminal kinesins. Proc. Natl. Acad. Sci. U. S. A. 108, 16253–16258 (2011).

37. McKenney, R. J., Huynh, W., Tanenbaum, M. E., Bhabha, G. & Vale, R. D. Activation of cytoplasmic dynein motility by dynactin-cargo adapter complexes. Science 345, (2014).

38. McKenney, R. J., Huynh, W., Vale, R. D. & Sirajuddin, M. Tyrosination of α‐tubulin controls the initiation of processive dynein–dynactin motility. EMBO J. 35, 1175–1185 (2016).

39. Ortega-Arroyo, J. & Kukura, P. Interferometric scattering microscopy (iSCAT): New frontiers in ultrafast and ultrasensitive optical microscopy. Phys. Chem. Chem. Phys. 14, 15625–15636 (2012).

40. Howard, J. Mechanics of Motor Proteins and the Cytoskeleton. (2001).

41. Schnitzer, M. J., Visscher, K. & Block, S. M. Force production by single kinesin motors. Nat. Cell Biol. 2, 718–723 (2000).

42. Cianfrocco, M. A., DeSantis, M. E., Leschziner, A. E. & Reck-Peterson, S. L. Mechanism and Regulation of Cytoplasmic Dynein. Annu. Rev. Cell Dev. Biol. 31, 83–108 (2015).

43. Urnavicius, L. et al. The structure of the dynactin complex and its interaction with dynein. Science 347, 1441–1446 (2015).

44. Elshenawy, M. M. et al. Lis1 activates dynein motility by pairing it with dynactin. bioRxiv 1–23 (2019). doi:https://doi.org/10.1101/685826

45. Htet, Z. M., Gillies, J. P., Baker, R. W. & Leschziner, A. E. Lis1 promotes the formation of maximally activated cytoplasmic dynein-1 complexes. bioRxiv 1–30 (2019). doi:10.1101/683052

46. Reck-Peterson, S. L., Redwine, W. B., Vale, R. D. & Carter, A. P. The cytoplasmic dynein transport machinery and its many cargoes. Nat. Rev. Mol. Cell Biol. 19, 382–398 (2018).

