## Supplementary Information for "Dynactin p150 promotes processive motility of DDB complexes by minimizing diffusional behavior of dynein"

**Supplement to Figure 5**

**State-switch detection algorithm**

Trajectories of x-y particle positions over time generated by FIESTA analysis software^1^ (**Fig. S1A**) were separated into displacements parallel and perpendicular to the microtubule by fitting a line to define the microtubule axis and rotating the trace (**Fig. S1 B**). If the particle position could not be fit by FIESTA for 10 consecutive frames (100 msec) due to background noise, the particle being out of focus, or other issues, the trace was terminated. The exception was if the particle was stuck in place during the dropped frames and could unambiguously be connected to a later trace when tracking resumed. If less than 10 consecutive frames were dropped, missing points were filled in by averaging previous and subsequent tracked positions.

For identifying motility states, a sliding 10-frame window was chosen for every time point (except the first 4 points and last 5 points of the trace), and three metrics were calculated: Velocity (V), standard deviation of position (SD_pos_), and standard deviation of the residual (SD_res_), defined as the SD_pos_ after subtracting the velocity. By examining many individual traces, three characteristic motility states were defined, as follows: Processive states (**Fig. S1C**) had high velocities and small SD_res_; Diffusive segments (**Fig. S1D**) had high SD_pos_ and/or high SD_res_; and Stuck segments (**Fig. S1E**) had low velocities and low SD_pos_. Thus, a motility state (P, D, or S) was defined for each time point (the 5^th^ point in a 10-frame window) using three cutoffs: V_cut_, SD_pos_cut_, and SD_res_cut_, as follows:

Processive state (P):	V > V_cut_	and	SD_res_ < SD_res_cut_

Stuck state (S):	V < V_cut_	and	SD_pos_ < SD_pos_cut_

Diffusive state (D)	SD > SD_pos_cut_	or	SD_res_ > SD_res_cut_

This initial cutoff-based analysis had two problems. First, it led to premature state-switch calls, because switching from processive or stuck states led to premature increases in SD_pos_ and/or SD_res_ at the corners. Second, transient fluctuations led to false-positive switch calls. These problems were solved by making a rule that a state-switch is called only when 5 consecutive time points (half of the 10-frame window) are called in the new state, and making the minimum state duration to be 10 points (100 msec). This lower duration limit is equivalent to limiting the switching rate to 10 s^-1^, which is reasonable for our 100 Hz data. This analysis resulted in the trace being broken into segments of defined states. From these data, the mean duration of each motility state, the fraction of time in spent each state, and the first-order switching rates between states were calculated.

**Data simulation**

For our initial guesses, we took the cutoffs as one standard deviation from the mean of the velocity distributions of stuck segments, SD_res_ distribution of processive segments and SD_pos_ distribution of stuck segments in **Fig. S1 F-H**; V_cut_ = 151 nm/s, SD_pos_cut_ = 12 nm, and SD_res_cut_ = 12 nm. The initial cutoffs used to assign the three motility states were chosen subjectively, so to refine the cutoffs, we carried out simulations with defined switches between states and analyzed the simulated data with our state detection algorithm. The first step in the simulations was to identify clear processive, diffusive and stuck episodes in our traces that were a minimum of 0.5 s long. We chose 10 P segments (17.3 s total), 5 D segments (3.2 s total), and 11 S segments (7.5 s total); examples are shown in **Fig. S1 C-E**. For each trace, we calculated V, SD_pos_ and SD_res_ for every timepoint using a 10-frame sliding window and analyzed their distributions (**Fig. S1 F – H)**. Particular features are apparent, such as the wide velocity distributions for processive and stuck segments (**Fig. S1F**), which result in part because they are calculated over only 100 msec intervals, as well as the fact that the stuck and processive velocity distributions overlap significantly. Secondly, there are reasonable separations of the SD_pos_ distributions for stuck and diffusive states (**Fig. S1E**) and the SD_res_ distributions for processive and diffusive states (**Fig. S1F**). Although it is possible to define cutoffs by eye from these distributions, we chose to identify optimum cutoffs using simulations and optimizing the algorithm to the simulated data with known state transition points.

To simulate P segments, segment velocities were chosen by sampling from the velocity distribution of processive segments in **Fig. S1 F**. Fluctuations around that mean velocity, which correspond to actual velocity fluctuations as well as experimental error and fitting uncertainty, were simulated by adding a white noise term that sampled from the SD_res_ distribution for processive segments in **Fig. S1 H**. For simulating S segments, velocity was set to zero, and positional variation was accounted for by adding a white noise term with a SD sampled from the stuck state SD_pos_ distribution shown in **Fig. S1 G**. Finally, D states were modeled by computing an effective 1D diffusion constant from mean-squared displacement analysis of the diffusive traces (**Fig. S1D inset**). To confirm the validity of how each state was simulated, we compared the distribution of 10-frame calculated V, SD_pos_ and SD_res_ from the simulations to those of the experimental data (**Fig. S2 D-F**).

Simulations were performed by starting motors in the S state, setting the mean state transition rate to 1 s^-1^ for every transition, choosing an exponentially distributed random number for the switch time to the two possible states (P and D in this case), and transitioning to the state with the smaller switching time. This process was repeated to generate a 500 s trace containing many transitions between motility states. Note that because each state could be exited by two possible transitions, the mean transition rate out of each state was 2 s^-1^, corresponding to a 0.5 s mean state duration time for each state in the simulations.

**Parameter optimization**

A key determinant of the accuracy of the switch detection algorithm is the choice of the cutoff parameters used to delineate the different states, V_cut_, SD_pos_cut_ and SD_res_cut_. To optimize our cutoff parameter choices, we applied our algorithm to the simulated data (**Fig. S2A**), quantified how well the algorithm correctly identified the known motility states (**Fig. S2 B, C**), and iteratively adjusted the cutoff parameters to achieve optimal performance of the algorithm. For a given cutoff parameter set, the performance of the algorithm took into account both the overall fraction of time the correct state was identified, as well as the relative error in calculating the duration of each motility state. The goodness of fit (GOF) parameter (where 1.0 is defined as a perfect fit) was calculated as:

$$GOF=0.5*\left( State accuracy \right)+ 0.5*\frac{{Dur}_{fit}^{P}+{Dur}_{fit}^{D}+{Dur}_{fit}^{S}}{3}$$

Here, *State accuracy* is the proportion of time points where the state was correctly called (**Fig. S2C**), and the accuracy of mean state durations is defined as:

$$Dur_{fit}= 1-\frac{\left| Detected duration-True duration \right|}{True duration}$$

The first problem we encountered was that V_cut_, used to delineate processive from stuck states, was poorly constrained. This issue can be seen in the overlap of Processive and Stuck velocities in **Fig. S1 F**, and means that any hard cutoff will miscall some segments. The mechanistic origin of broad distribution of DDB velocities, which has consistently been observed by others in both raw kymographs and in velocity distribution plots ^2–6^, is not understood. The relationships between the P and S velocity distributions can be seen more clearly when plotted as cumulative distributions (**Fig. S3 A**). Based on this visual analysis, we chose a compromise V_cut_ of 100 nm/s, which results in a nearly equal balance of 78% of processive states and 81% of stuck states being correctly called.

To identify the optimal SD_pos_cut_ and SD_res_cut_ parameters, these two parameters were varied while holding V_cut_ constant at 100 nm/s, and for each parameter set, a goodness of fit (GOF) value was calculated. A GOF heat map (**Fig. S3 B**) shows that a GOF maximum of 80% was identified for SD_pos_cut_ = 10 nm and SD_res_cut_ = 18 nm. These parameter choices were used in all subsequent analysis of the experimental data. To confirm that the 100 nm/s velocity cutoff (chosen based on **Fig. S3A**) was consistent with the GOF optimization, the GOF optimization was repeated while varying all three parameters. As can be seen in **Fig. S3 C, D**, for any SD_pos_cut_ or SD_res_cut_ parameter choice along the x-axis, changing V_cut_ along the y-axis produced minimal color changes. Specifically, increasing V_cut_ above 100 nm/s led to negligible improvements in the GOF (all GOF > 80%), validating our V_cut_ choice of 100 nm/s. However, it should be noted that even with this cutoff, it means that segments where DDB is moving steadily at below 100 nm/s are defined as stuck states. The molecular mechanism underlying these slow-moving states is not known. We hypothesize that either the motors are partially active or else there is a non-motor portion of DDB that binds tightly to the microtubule under some conditions, resulting in long-duration slow motility. To conclude, from the simulation-based parameter optimization, the final cutoff values were: V_cut_ = 100 nm/s, SD_res_cut_ = 18 nm, and SD_pos_cut_ = 10 nm.

**Sensitivity analysis for experimental data**

To assess how sensitive the results are to the specific choices of parameters, we varied the cutoff values up and down by 20% and calculated the changes in the mean state durations and fraction of time spent each state. As shown in **Fig. S4 and S5**, the state durations and state fractions did vary somewhat with changes in the cutoff parameters. For instance, changing V_cut_ led to reciprocal changes in mean durations of Processive and Stuck segments (**Fig. S4 A and G**) and in the fraction of time in Processive and Stuck states (**Fig. S5 A and G**). However, in all cases, the differences between durations and state fractions between DDB control and DDB in the presence of p150 antibody were robust against 20% changes in parameters. Thus, the qualitative differences identified between DDB control and in the presence of Ab_p150_ are not dependent on the specific choices used for the cutoff parameters.

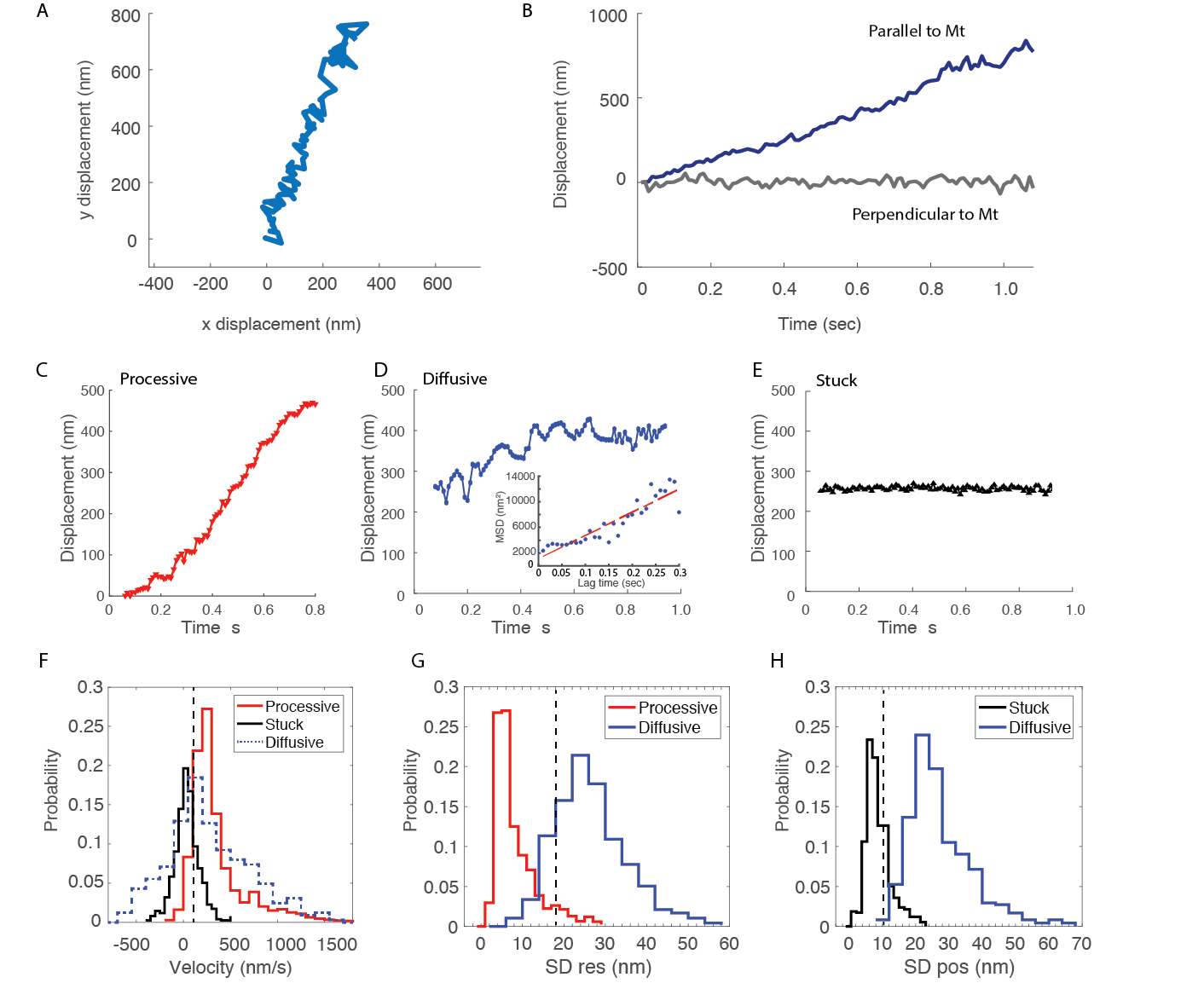

**Figure S1: Data processing and characteristics of processive, diffusive and stuck states**

**(A)** Sample trace of x-y position of particle-tagged DDB. **(B)** Time-dependent displacement parallel (blue) and perpendicular (black) to the microtubule (Mt). **(C-E)** Representative traces of processive, diffusive and stuck episodes. **D inset:** Fig of mean-squared displacement for the diffusive episode, resulting in a calculated diffusion constant, D = 20,000 nm^2^/s. **(F)** Distribution of velocities from 10-frame (100 ms) windows for processive (red), stuck (black) and diffusive (blue dashed) segments. **(G)** Distribution of residual standard deviation following subtraction of slope for processive (red) and diffusive (blue) segments. **(H)** Distribution of positional standard deviation for stuck (black) and diffusive (blue) segments. For all distributions, V, SD_pos_ and SD_res_ values were calculated over 10-frame windows from 100 frame/s videos. Dashed lines indicate parameter cutoff choices based on simulation analysis described below.

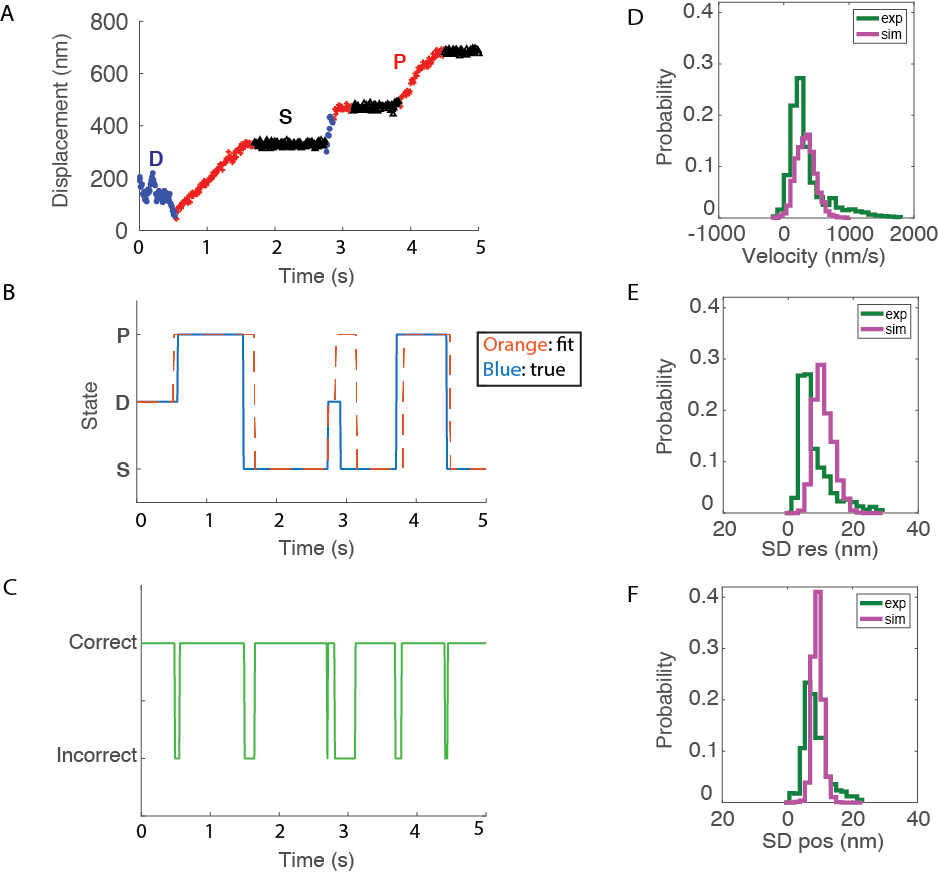

**Figure S2: Simulations used to refine switch detection algorithm**

**(A)** Simulated trace with states identified by the switch point detection algorithm using initial guesses for cutoff parameters. **(B)** Corresponding plot of actual (blue) and algorithm-identified (orange) states. **(C)** Corresponding plot of correct versus incorrect state identification using this cutoff parameter set. **(D, E, F)** Comparison of distributions for 10-frame running windows for simulated data (purple) and experimental data (green; taken from **Fig. S1F-H**) for processive segment velocity (**D**), processive segment SD_res_ (**E**) and stuck segment SD_pos_ (**F**).

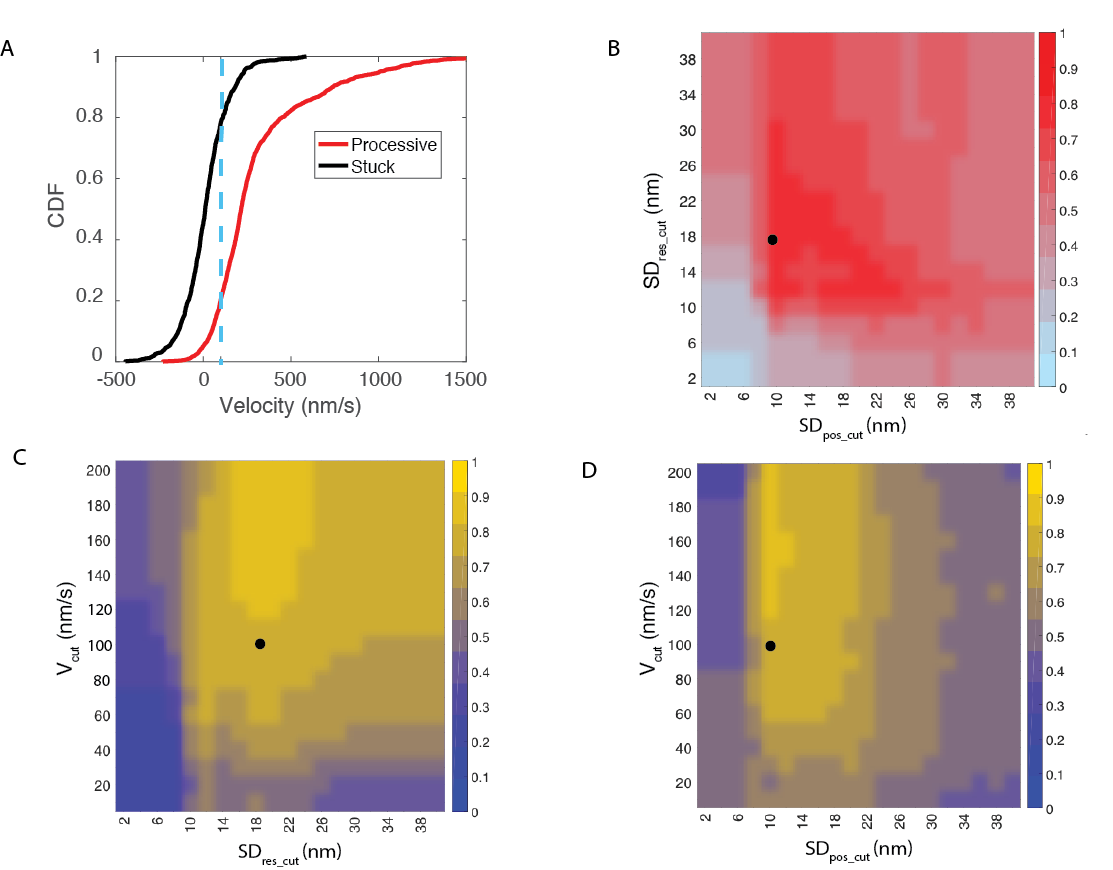

**Figure S3: Parameter optimization and comparison of segment velocity distributions**

**(A)** Cumulative distributions of stuck (black) and processive (red) segment velocities from experimental data (replotted from distributions in **Fig. S1 F**). Blue line denotes V_cut_ of 100 nm/s where 78% of stuck velocities are below and 81 % of processive velocities are above the cutoff. **(B)** Heat map of goodness of fit (GOF) parameter from algorithm analysis of simulated data. V_cut_ was set to 100 nm/s, and performance shown for different SD_pos_cut_ and SD_res_cut_ values. Optimal algorithm detection results (GOF = 80 %) were obtained at SD_pos_cut_ = 10 nm and SD_res_cut_ =18 nm (denoted by black dot). **(C)** GOF heat map to simulated data using algorithm with SD_res_cut_ fixed at 18 nm and varying V_cut_ and SD_pos_cut_. Optimal results (GOF> 80%) were obtained when SD_pos_cut_ = 10 nm and V_cut_ > 100 nm/s (black dot). **(D)** GOF heat map to simulated data using algorithm with SD_pos_ fixed at 10 nm and varying V_cut_ and SD_res_cut_. Optimal results (GOF> 80%) were obtained when SD_Res_cut_ = 18 nm and V_cut_ > 100 nm/s (black dot). Based on this analysis, the cutoff parameters for subsequent analysis were: V_cut_ = 100 nm/s, SD_pos_cut_ = 10 nm, and SD_res_cut_ = 18 nm.

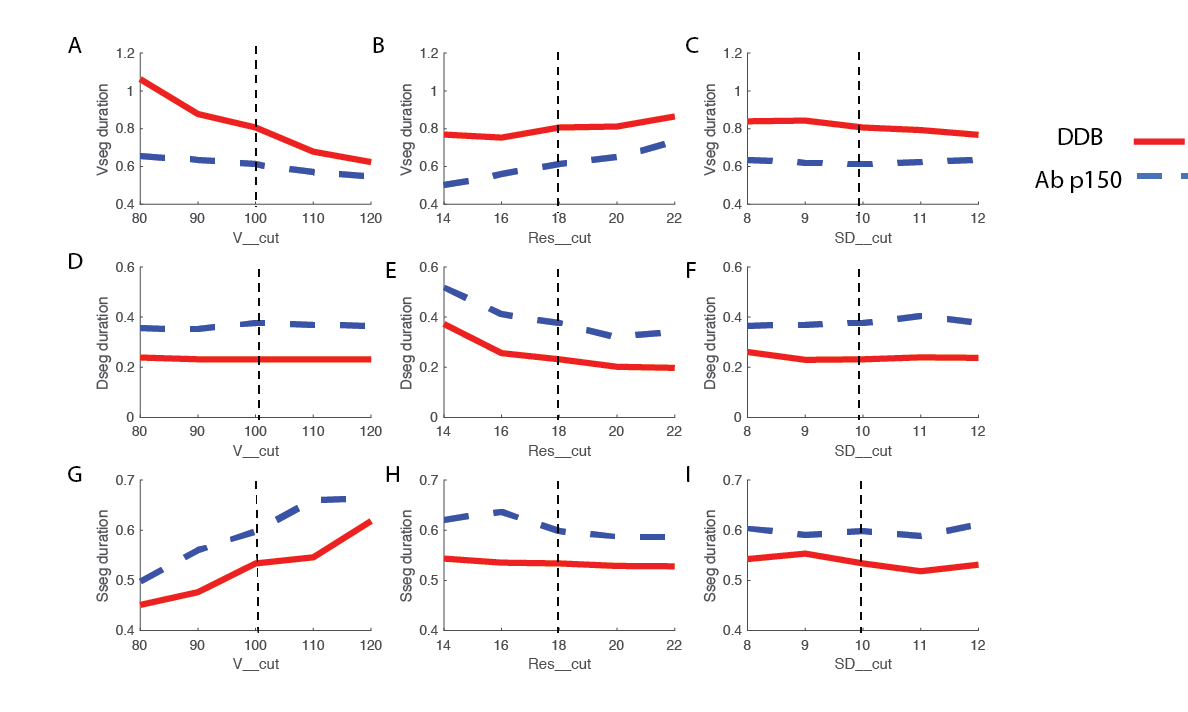

**Figure S4: Cutoff parameter sensitivity analysis for identifying segment durations in experimental data.** Each plot shows mean P, D and S segment durations for DDB control (red solid lines) and Ab_p150_ (blue dashed lines) for the simulation-optimized cutoff values (vertical dashed lines) and variations of each by +/- 20%. Importantly, although the precise estimates of the mean durations change somewhat with changes in the cutoff parameters, the relative difference between durations for control and Ab_p150_ groups remains relatively constant. Thus, the segment duration results are robust (not strongly sensitive) with regard to the cutoff parameter choices.

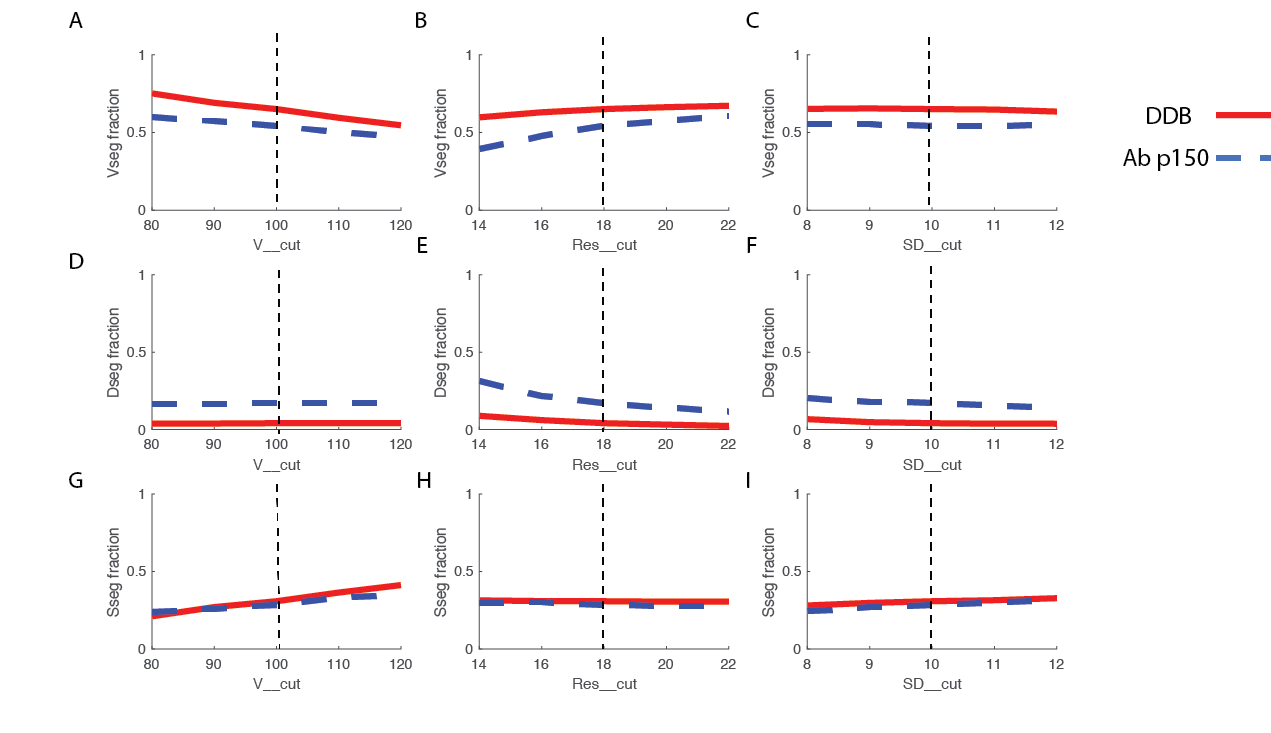

**Figure S5: Cutoff parameter sensitivity analysis for identifying fraction of time spent in each state in experimental data.** Each plot shows fraction of time spent in P, D or S states for DDB control (red solid lines) and Ab_p150_ (blue dashed lines) for the simulation-optimized cutoff values (vertical dashed lines) and variations of each by +/- 20%. Importantly, although the precise estimates of the fractions of time spent in each state change somewhat with changes in the cutoff parameters, the relative differences between control and Ab_p150_ groups remains relatively constant. Thus, results for the fraction of time spent in each state are robust (not strongly sensitive) with regard to the cutoff parameter choices.

**References:**

1.	Ruhnow, F., Zwicker, D. & Diez, S. Tracking Single Particles and Elongated Filaments with Nanometer Precision. *Biophysj* **100**, 2820–2828 (2011).

2.	McKenney, R. J., Huynh, W., Tanenbaum, M. E., Bhabha, G. & Vale, R. D. Activation of cytoplasmic dynein motility by dynactin-cargo adapter complexes. *Science (80-. ).* **345**, (2014).

3.	Belyy, V. *et al.* The mammalian dynein/dynactin complex is a strong opponent to kinesin in a tug-of-war competition. **18**, 1018–1024 (2017).

4.	Urnavicius, L. *et al.* Cryo-EM shows how dynactin recruits two dyneins for faster movement. *Nature* **554**, 202–206 (2018).

5.	Grotjahn, D. A. *et al.* Cryo-electron tomography reveals that dynactin recruits a team of dyneins for processive motility. *Nat. Struct. Mol. Biol.* **25**, 203–207 (2018).

6.	Gutierrez, P. A., Ackermann, B. E., Vershinin, M. & Mckenney, R. J. Differential effects of the dynein-regulatory factor Lissencephaly-1 on processive dynein-dynactin motility. *J. Biol. Chem.* **292**, 12245–12255 (2017).

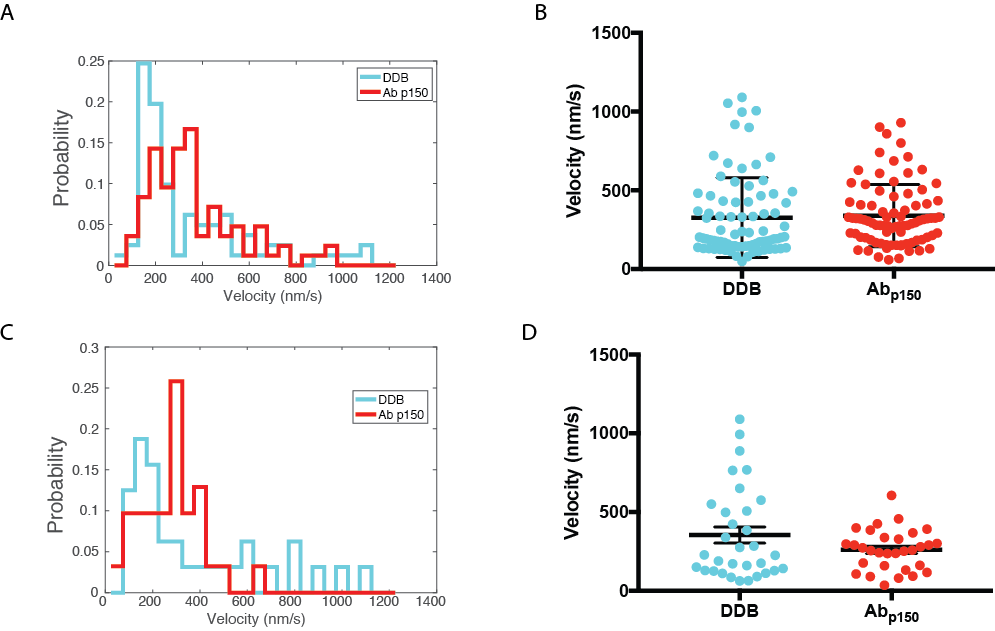

**Figure S6 (Related to Figure 6): Velocity distributions.**

**(A)** Distribution of velocities of processive segments identified by the detection algorithm for DDB control (blue, n=81) and Ab_p150_ (red, n=84). Velocities were calculated by linear regression to each processive segment. **(B)** Comparison of mean processive segment velocities, showing no statistical difference (n.s., p =0.72 using two-tail T-test) between DDB control (blue) and Ab_p150_ (red). **(C)** Distribution of whole-trace velocities for DDB control (blue, n=32) and Ab_p150_ (red, n=31). Velocities were calculated by linear regression to entire traces. **(D)** Comparison of mean whole-trace velocities, showing no statistical difference (n.s., p =0.104 using two-tail T-test) between DDB control (blue) and Ab_p150_ (red).

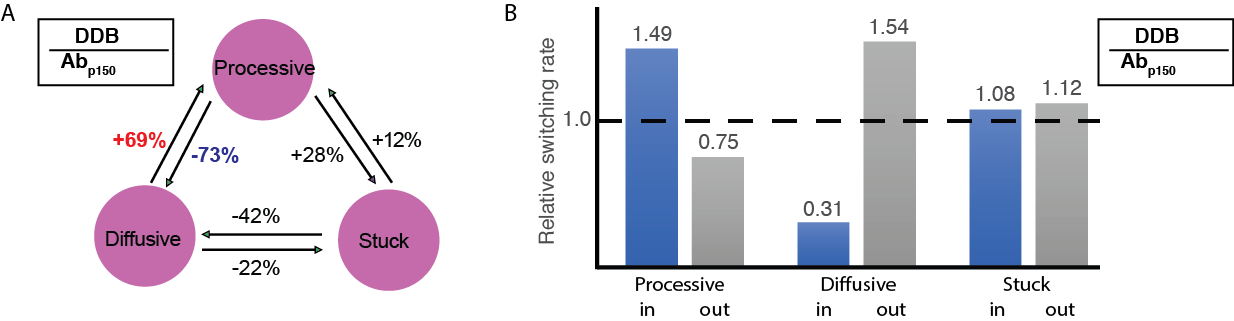

**Figure S7 (Related to Figure 6): Relative changes in switching rates.**

**(A)** Relative changes in switching rates when dynactin p150 is able to interact with microtubules. The prominent increase in the switching rate from diffusive to processive is denoted by red, and the prominent decrease in switching from diffusive to processive is denoted by blue. **(B)** Relative changes in overall switching rates into and out of each state when dynactin p150 is blocked by antibody. Most notably, the presence of p150 causes an increase in switching into the processive state and a decrease in switching out of the processive state. These changes primarily result from changes into and out of the diffusive state and not to changes into or out of the stuck state.

**Video S1: Motility of gold-labeled DDB by iSCAT.**

DDB complexes functionalized with 30-nm gold nanoparticles were imaged moving along immobilized microtubules at 100 frames/s by iSCAT microscopy. Movies were processed by subtracting still images of microtubule and inverting to produce bright particle on dark background.

**Video S2: Bidirectional transport of kinesin-1 – DDB complexes by TIRF.**

Images at right are Qdot channel, showing kinesin-DDB complexes moving slowly and bidirectionally along microtubules. Images at left are from GFP channel, showing streaming of excess free kinesin-1 motors to the microtubule plus-end; this information is used to assign microtubule polarity. Qdot and GFP movies were taken sequentially. Arrows in first frame denote: cargos transported to the plus end (yellow arrow), cargos transported to the minus end (red arrow), and cargos that switch from plus-end to minus-end direction (white arrow).
